# CRISPR RNA-guided integrases for high-efficiency and multiplexed bacterial genome engineering

**DOI:** 10.1101/2020.07.17.209452

**Authors:** Phuc Leo H. Vo, Carlotta Ronda, Sanne E. Klompe, Ethan E. Chen, Christopher Acree, Harris H. Wang, Samuel H. Sternberg

## Abstract

Tn*7*-like transposons are pervasive mobile genetic elements in bacteria that mobilize using heteromeric transposase complexes comprising distinct targeting modules. We recently described a Tn*7*-like transposon from *Vibrio cholerae* that employs a Type I-F CRISPR–Cas system for RNA-guided transposition, in which Cascade directly recruits transposition proteins to integrate donor DNA downstream of genomic target sites complementary to CRISPR RNA. However, the requirement for multiple expression vectors and low overall integration efficiencies, particularly for large genetic payloads, hindered the practical utility of the transposon. Here, we present a significantly improved INTEGRATE (insertion of transposable elements by guide RNA-assisted targeting) system for targeted, multiplexed, and marker-free DNA integration of up to 10 kilobases at ~100% efficiency. Using multi-spacer CRISPR arrays, we achieved simultaneous multiplex insertions in three genomic loci, and facile multi-loci deletions when combining orthogonal integrases and recombinases. Finally, we demonstrated robust function in other biomedically- and industrially-relevant bacteria, and developed an accessible computational algorithm for guide RNA design. This work establishes INTEGRATE as a versatile and portable tool that enables multiplex and kilobase-scale genome engineering.

---

DNA technologies to stably integrate genes and pathways into the genome enable the generation of engineered cells with entirely new functions. Applications of this powerful approach have already yielded impactful commercial products, with examples including CAR-T cell therapies^1^, genetically modified crops^2^, and cell factories producing diverse compounds and medicines^3^. In many of these applications, genomic integration is highly preferred over plasmid-based methods for maintaining heterologous genes in engineered cells, due to improved stability in the genome, better control of copy numbers, and regulatory concerns regarding biocontainment of recombinant DNA^4,5^. However, generation of modified cells with kilobases of changes across the genome remains practically challenging, often requiring inefficient, multi-step processes that are time and resource intensive. A facile, efficient and versatile method that allows programmable genomic integrations in multiplex would significantly accelerate advances in cellular programming for many application areas.

In bacteria, genome engineering and integration can be achieved through several approaches that utilize endogenous or foreign integrases^4,5^, transposases^6,7^, recombinases^8,9^, or homologous recombination (HR) machinery^10–13^, which can be further combined with CRISPR-Cas to improve efficiency^14,15^. While widely used, these methods are not without significant drawbacks. For example, recombination-mediated genetic engineering (recombineering) using λ-red or RecET recombinase systems in *E. coli* allows programmable genomic integrations, specified by the homology arms flanking the foreign DNA cassette^13,16^. However, recombineering efficiency is generally low (less than 1 in 10^3^–10^4^)^17^ without selection of a co-integrating selectable marker^8^ or CRISPR-Cas-mediated counter-selection of unedited alleles^14^, and thus cannot be easily multiplexed to make simultaneous insertions into the same cell. There is a limited number of robust selectable markers (e.g. antibiotic resistance genes) that require another excision step to remove from the genome for subsequent reuse, and expression of Cas9 for negative selection can cause unintended DNA double-strand breaks (DSBs) that lead to cytotoxicity^18–20^. Practically, recombineering has a payload size limit of only 3-4 kb in many cases, making it less useful for genomic integration of pathway-sized DNA cassettes. Finally, unknown requirements for host-specific factors or cross-species incompatibilities of phage recombination proteins have rendered *E. coli* recombineering systems more challenging to port to other bacteria, requiring significant species-specific optimizations^21^ or screening of new recombinases^22^.

Other integrases and transposases, such as ICEBs1 and Tn7, have also been used for genome integration^4,23^. These systems recognize highly specific attachment sites that are unfortunately difficult to reprogram, and thus require the prior presence of these sites or their separate introduction in the genome^24^. Other more portable transposons, such as Mariner and Tn5, generate non-specific integrations that have been used for genome-wide transposon mutagenesis libraries^25–28^. However, these transposases cannot be targeted to specific genomic loci, and large-scale screens are needed to isolate desired clones. More recently, a catalytically-dead Cas9 has been fused to either a transposase or a recombinase to provide better site specificity, which showed success in mostly *in vitro* studies^29,30^. Autocatalytic Group II RNA introns, selfish genetic elements in bacteria, have also been used for genomic transpositions and insertions^31,32^. This system utilizes an RNA intermediate to guide insertions, but suffers from inconsistent efficiencies ranging from 1-80% depending on the target site and species^33^, and a limited cargo size of 1.8 kb^34^.

An ideal genome insertion technology should provide efficient single-step DNA integration for high capacity cargos with high specificity and programmability, without relying on DSBs and HR. We recently described a new category of programmable integrases whose sequence specificity is governed exclusively by guide RNAs^35^. Motivated by the bioinformatic description of Tn*7*-like transposons encoding nuclease-deficient CRISPR-Cas systems^36^, we selected a candidate CRISPR-transposon from *Vibrio cholerae* (Tn*6677*) and reconstituted RNA-guided transposition in an *E. coli* host. DNA integration occurs ~47–51 base pairs (bp) downstream of the genomic site targeted by the CRISPR RNA (crRNA), and requires transposition proteins TnsA, TnsB, and TnsC, in conjunction with the RNA-guided DNA targeting complex TniQ-Cascade^35,37^ (**Fig. 1a,b**). Remarkably, bacterial transposons have hijacked at least three distinct CRISPR–Cas subtypes^35,38,39^, and work from Zhang and colleagues demonstrated that the Type V-K effector protein, Cas12k, also directs targeted DNA integration, albeit with lower fidelity^40^. These studies underscore the exaptation that allowed transposons to repurpose RNA-guided DNA targeting systems for selfish propagation, and highlight the promise of programmable integrases, which we coined INTEGRATE (insertion of transposable elements by guide RNA-assisted targeting), for genome engineering^41,42^.

**Fig. 1.**
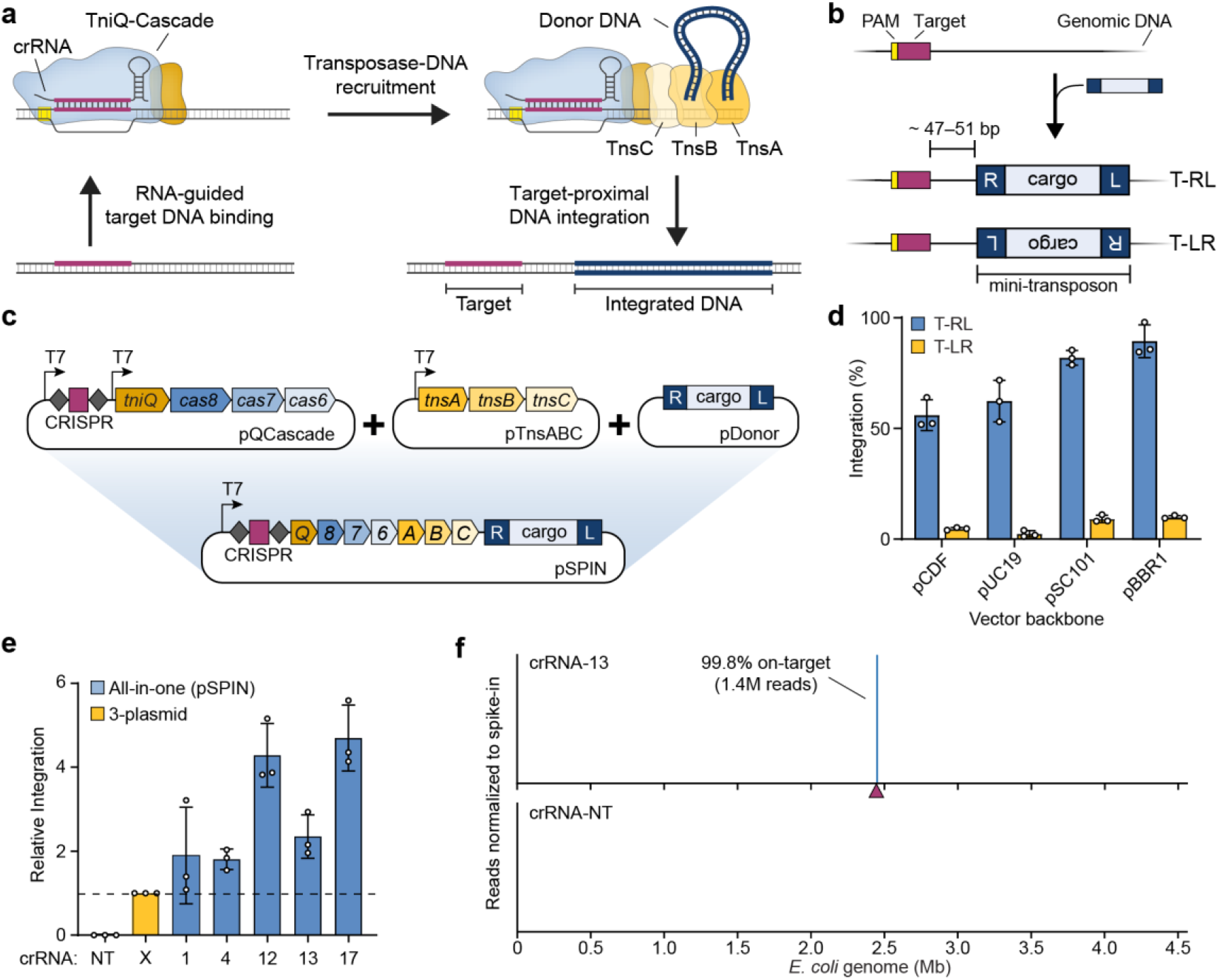
Streamlined single-plasmid system for RNA-guided DNA integration. **a,** Schematic of INTEGRATE (insertion of transposable elements by guide RNA-assisted targeting) using a *Vibrio cholerae* CRISPR-transposon. **b,** RNA-guided DNA integration occurs ~47-51 bp downstream of the target site, in one of two possible orientations (T-RL and T-LR); the donor DNA comprises a genetic cargo flanked by left (L) and right (R) transposon ends. **c,** Top, a three-plasmid INTEGRATE system encodes protein-RNA components on pQCascade and pTnsABC, and the donor DNA on pDonor. Bottom, a single-plasmid INTEGRATE system (pSPIN) drives protein-RNA expression with a single promoter, on the same vector as the donor DNA. **d,** qPCR-based quantification of integration efficiency with crRNA-4, for pSPIN containing distinct vector backbones of differing copy numbers. **e**, Relative integration efficiencies for the three-plasmid or single-plasmid (pSPIN) expression system across five distinct crRNAs. Data are normalized to the three-plasmid system; pSPIN contained the pBBR1 backbone. **f,** Normalized Tn-seq data for crRNA-13 and a non-targeting crRNA (crRNA-NT) for pSPIN containing the pBBR1 backbone. Genome-mapping reads are normalized to the reads from a spike-in control (**Methods**); the target site is denoted by a maroon triangle. Data in **d** and **e** are shown as mean ± s.d. for n = 3 biologically independent samples.

INTEGRATE benefits from both the high-efficiency, seamless integrations of transposases, as well as the simple programmability of CRISPR-mediated targeting. However, the system previously demonstrated in *E. coli* required multiple cumbersome genetic components and displayed low efficiency for larger insertions in dual orientations. Here, we developed a vastly improved INTEGRATE system that use streamlined expression vectors to direct highly accurate insertions at ~100% efficiency effectively in a single orientation, independent of the cargo size, without requiring selection markers. Since INTEGRATE does not rely on HR, multiple simultaneous genomic insertions into the same cell could be generated using CRISPR arrays with multiple targeting spacers, and INTEGRATE paired with Cre-Lox was used to achieve multiple genomic deletions. Finally, we demonstrated the portability and high-accuracy of INTEGRATE in other species, including *Klebsiella oxytoca* and *Pseudomonas putida*, highlighting its broad utility for bacterial genome engineering.

## RESULTS

### An optimized, single-plasmid system for high-efficiency RNA-guided DNA integration

We previously employed a three-plasmid expression system to reconstitute RNA-guided DNA integration in *E. coli*, whereby pQCascade and pTnsABC encoded the necessary protein-RNA components, and pDonor contained the mini-transposon (mini-Tn, aka donor DNA) (**Fig. 1c**). To streamline our strategy and eliminate both antibiotic burden and the need for multiple transformation events, we serially reduced the number of independent promoters and plasmids and ultimately arrived at a single-plasmid INTEGRATE construct (pSPIN), in which one promoter drives expression of the crRNA and polycistronic mRNA, directly upstream of the mini-Tn (**Fig. 1c**, **Supplementary Fig. 1**, and **Supplementary Table 1**). This design allows modular substitution of the promoter and/or genetic cargo for user-specific applications, and straightforward subcloning into distinct vector backbones.

After identifying a functionally optimal arrangement of the CRISPR array and operons (**Supplementary Fig. 1**), we transformed *E. coli* BL21(DE3) with four pSPIN derivatives encoding a *lacZ*-specific crRNA on distinct vector backbones, and monitored the efficiency of RNA-guided transposition by quantitative PCR (qPCR). Surprisingly, our streamlined plasmids exhibited enhanced integration activity, with efficiency exceeding 90% using the pBBR1 vector backbone (**Fig. 1d**), and showed substantially stronger bias for insertion events in which the transposon right end was proximal to the target site (T-RL), as compared to the original three-plasmid expression system (**Supplementary Fig. 2**). To determine whether increased efficiency would translate across multiple targets, we assessed integration at five target sites used in our previous study^35^ and found that our pSPIN vector was consistently 2–5X more efficient (**Fig. 1e**). Our single-plasmid INTEGRATE system maintained high-fidelity activity, and an absence of insertion events with a non-targeting crRNA, as reported by genome-wide transposon-insertion sequencing (Tn-seq; **Fig. 1f** and **Supplementary Fig. 3**). This high degree of specificity was further verified by isolating clones and confirming the unique presence of a single insertion by whole-genome, single-molecule real-time (SMRT) sequencing and structural variant analysis (**Methods** and **Supplementary Table 2**).

We next assessed the role of expression level by modifying the promoter driving protein-RNA expression. Using a panel of constitutive promoters of varying expression strength, we found that higher expression drove higher rates of integration, without any deleterious effect on genome-wide specificity (**Fig. 2a,b** and **Supplementary Fig. 4a**). Efficient integration was also driven by a natural broad-host promoter recently adopted for metagenomic microbiome engineering^43^ (**Fig. 2a**), and the use of constitutive promoters allowed us to demonstrate high-accuracy integration in additional *E. coli* strains, including MG1655 and BW25113, without any requirement for host recombination factors (**Supplementary Fig. 4b,c**). Interestingly, we also noticed that RNA-guided DNA integration readily proceeded when cells were grown at room temperature, and reached ~100% efficiency (without selection for the integration event) while maintaining 99.7% on-target specificity, even for the low-strength J23114 promoter (**Fig. 2c** and **Supplementary Fig. 4d**). This temperature-dependent activity may be due to the slower doubling time allowing for a greater window of opportunity, similar to a recent study of CRISPR–Cas activity in *Pseudomonas aeruginosa*^44^, or may reflect the natural operating range of enzymes derived from *V. cholerae*.

**Fig. 2.**
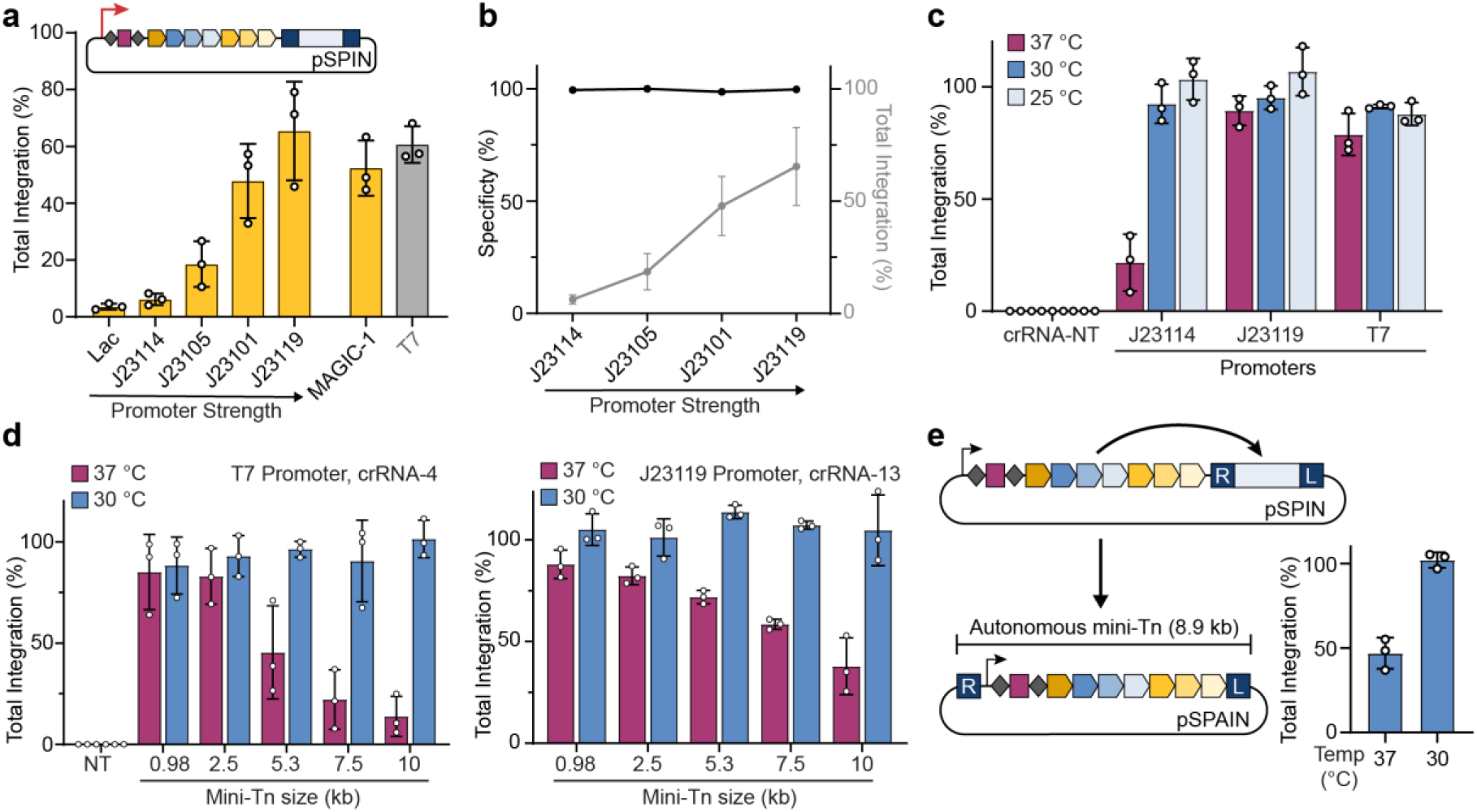
INTEGRATE supports high-efficiency insertion of large (10-kb) genetic payloads. **a,** qPCR-based quantification of integration efficiency with crRNA-4 as a function of pSPIN promoter identity. **b,** DNA integration specificity (black) for the promoters shown, as determined by Tn-seq, calculated as the percent of on-target reads relative to all genome-mapping reads (**Methods**); total integration efficiencies (qPCR) are plotted in grey. **c,** qPCR-based quantification of integration efficiency for crRNA-4 as a function of culture temperature and promoter strength. Integration reaches ~100% efficiency at lower growth temperatures for all constructs, including the weaker J23114 promoter. **d,** qPCR-based quantification of integration efficiency for variable mini-Tn sizes after culturing at either 30 or 37 °C. The promoter and crRNA used in each panel are shown at top; experiments were performed with a two-plasmid system comprising pEffector (pEffector-B, **Supplementary Fig. 1c**) and pDonor. Unless specified, transposition assays elsewhere in this study use a 0.98-kb mini-Tn. **e,** qPCR-based quantification of integration efficiency with crRNA-4 for a single-plasmid autonomous INTEGRATE system (pSPAIN), after culturing at 30 and 37 °C. The inserted DNA encodes all the necessary machinery for further mobilization. Integration efficiency data in **a-e** are shown as mean ± s.d. for n = 3 biologically independent samples.

Motivated by this finding, we reexamined the effect of cargo size on integration efficiency. We previously found that while the *V. cholerae* machinery integrated a ~1-kb cargo with optimal efficiency, larger cargos were poorly mobilized^35^. Remarkably, when we expressed protein-RNA components from a single effector plasmid (pEffector-B, **Supplementary Fig. 1c**) and cultured cells at 30 °C, we could integrate mini-transposons spanning 1–10 kb with ~100% efficiency, with no observable size-dependent effects (**Fig. 2d**) and without the need for marker selection. The same pattern was observed across multiple target sites and promoters, and the specificity of 10-kb insertions was verified by Tn-seq and SMRT sequencing (**Fig. 2d** and **Supplementary Fig. 4e-g**). As native CRISPR-containing transposons are frequently several tens of kb in size^35,36^, we anticipate that the system has the potential to mobilize payloads beyond 10 kb. To further leverage the large-cargo capability, we generated a single-plasmid autonomous INTEGRATE system (pSPAIN) in which the protein-RNA coding genes were cloned within the mini-Tn itself, and showed that this construct also directed targeted integration at ~100% efficiency (**Fig. 2e**). We envision that autonomous INTEGRATE systems, by virtue of mobilizing themselves according to the user-defined CRISPR array content, could serve as potent gene drive elements capable of programmed self-propagation in mixed community environments.

### Development of orthogonal integrases for iterative DNA insertions

As strain engineering often requires multiple insertions and knockouts to be performed in distinct genomic regions, we next sought to evaluate the amenability of the *V. cholerae* system for iterative integration events. We first cloned a derivative of pSPIN using a temperature-sensitive plasmid backbone, isolated a clonal strain containing a *lacZ*-specific insertion (target-4), and cured the plasmid (**Methods**). Next, we re-introduced the machinery to generate a proximal insertion at variable distances upstream of target-4, but using a mini-Tn whose distinct cargo could be selectively tracked by qPCR (**Fig. 3a**). Previous studies have demonstrated that Tn*7* and Tn*7*-like transposons exhibit target immunity^40,45,46^, whereby integration is prevented at target sites already containing another transposon copy. When we compared integration across a panel of crRNAs for strains with and without a pre-existing mini-Tn, we found that the *V. cholerae* transposon also exhibits target immunity, with ~20% relative efficiency at target sites ~5-kb away (**Fig. 3a** and **Methods**). This effect was ablated when we instead targeted a *glmS*-proximal site (target-1) that was >1 Mbp from the pre-existing insertion, demonstrating that iterative insertions are straightforward but more efficient when spaced far apart.

**Fig. 3.**
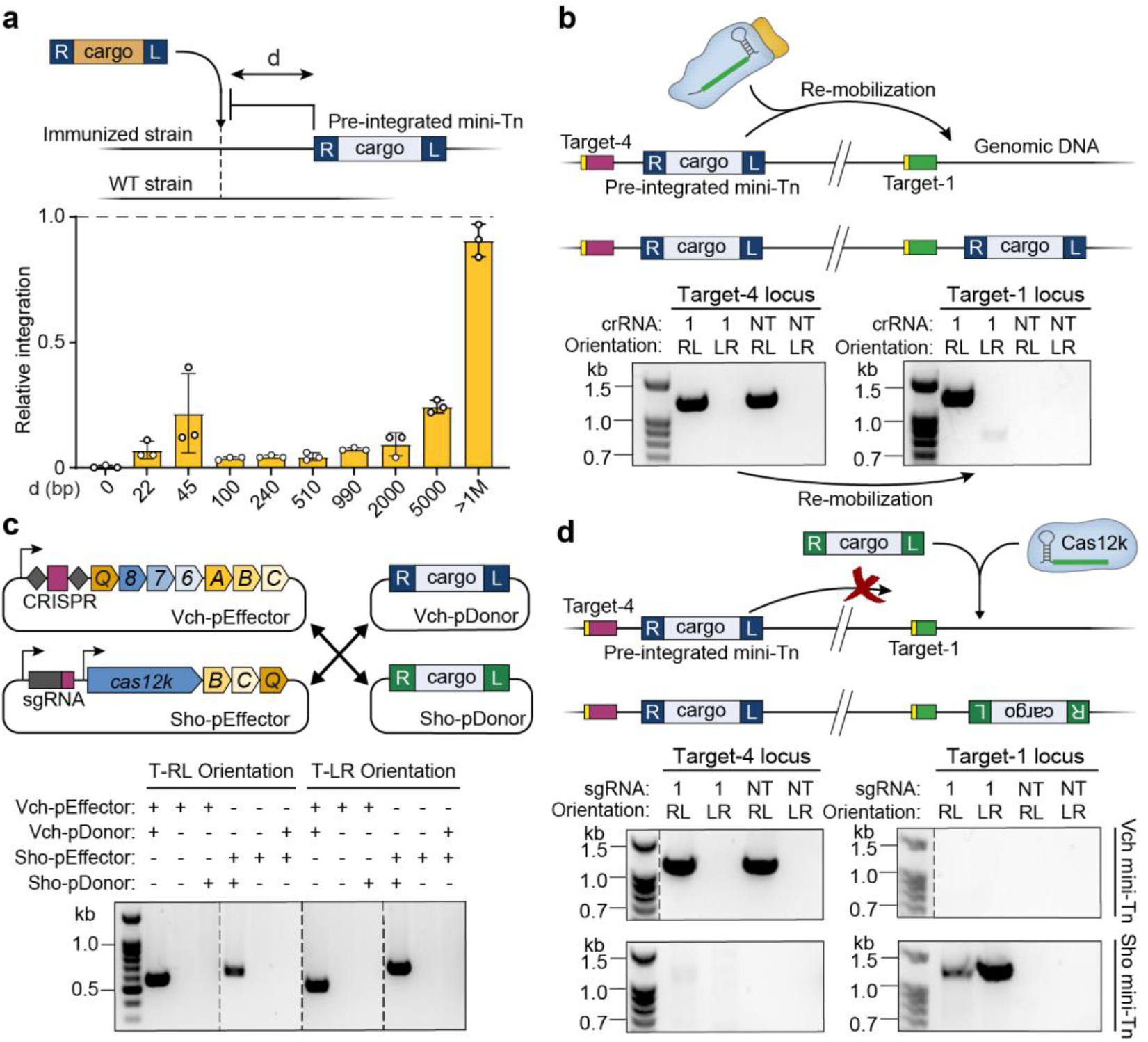
Orthogonal INTEGRATE systems facilitate multiple, iterative insertions. **a,** Effect of target immunity on RNA-guided DNA integration. An *E. coli* strain containing a single genomically integrated mini-Tn was generated, and the efficiency of additional transposition events using crRNAs targeting d bp upstream was determined by qPCR. Plotted is the relative efficiency for each crRNA in the immunized versus wild-type strain. **b,** Top, schematic showing re-mobilization of a genomically integrated mini-Tn (target-4) to a new genomic site (target-1) with crRNA-1. Bottom, PCR products probing for the mini-Tn at target-4 (left) and target-1 (right), resolved by agarose gel electrophoresis. The mini-Tn is efficiently transposed to target-1 by crRNA-1, without apparent loss of the mini-Tn at target-4. **c,** Top, schematic of orthogonal INTEGRATE systems from *V. cholerae* (Vch; Type I-F) and *S. hofmannii* (Sho; Type V-K), in which pDonor is separate from pEffector. Bottom, PCR products probing for RNA-guided DNA integration at target-4 with both systems, resolved by gel electrophoresis. Integration only proceeds with a cognate pairing between the expression and donor plasmids. **d,** Top, schematic of a second DNA insertion made by leveraging the orthogonal ShoINT system, for which the Vch mini-Tn is inert. Bottom, PCR products probing for either the Vch mini-Tn (top) or Sho mini-Tn (bottom) at target-4 (left) and target-1 (right), resolved by agarose gel electrophoresis. The Sho mini-Tn is efficiently integrated at target-1 by sgRNA-1, without loss of the Vch mini-Tn at target-4. Data in **a** are shown as mean ± s.d. for n = 3 biologically independent samples.

The simultaneous presence of a genomically integrated mini-Tn and distinct plasmid-borne mini-Tn produces an interesting scenario in which the transposase machinery can theoretically employ either DNA molecule as the donor substrate for integration (**Supplementary Fig. 5a**). Using cargo-specific primers, we showed that new insertions at target-1 were indeed a heterogeneous mixture of both mini-Tn donors, although the higher-copy plasmid source was heavily preferred (**Supplementary Fig. 5a**). To further investigate intramolecular transposition events, we transformed our clonally integrated strain with a plasmid encoding the protein-RNA machinery without donor DNA, and monitored re-mobilization of the pre-existing mini-Tn from target-4 to target-1. Integration at target-1 was readily observed, but surprisingly, we saw no PCR evidence of mini-Tn loss at target-4, despite the expectation that the transposon mobilizes through a cut-and-paste mechanism^35^ (**Fig. 3b** and **Methods**), suggesting that lesions resulting from donor DNA excision are rapidly resolved by HR, as has been observed with Tn*7*^47^.

To avoid any low-level contamination between donor DNA molecules, we explored the use of multiple RNA-guided transposases whose cognate transposon ends would be recognized orthogonally. Guided by prior bioinformatic description and experimental validation of transposons encoding Type V-K CRISPR–Cas systems^35,38,40^, we developed a new INTEGRATE system derived from *Scytonema hofmannii* strain PCC 7110 (hereafter ShoINT, **Supplementary Fig. 5b**); we note that the protein components are 30–55% identical to the homologous system described by Strecker and colleagues (ShCAST)^40^, which derives from a distinct *S. hofmannii* strain (**Supplementary Table 3**). ShoINT catalyzes RNA-guided DNA integration with 20–40% efficiency, and strongly favors integration in the T-LR orientation, albeit with detectable bidirectional integration at multiple target sites (**Supplementary Fig. 5c–e**). Next, we combined pEffector plasmids for the *V. cholerae* INTEGRATE system (VchINT) or ShoINT with either its own cognate pDonor or pDonor from the other system, and found that each RNA-guided integrase was exclusively active on its respective mini-Tn substrate (**Fig. 3c**). With this knowledge, we were able to sequentially introduce a new cargo at different locus (at target-1) using ShoINT, without any secondary mobilization of the pre-existing VchINT mini-Tn (at target-4) (**Fig. 3d)**; we expect the same lack of cross-reactivity between VchINT and ShCAST, based on similarities between the Type V-K proteins and transposon ends. This approach of using systems with transposon ends that are sufficiently distinct will enable orthogonal and iterative integration events for distinct genetic payloads.

We were keen to carefully compare genome-wide integration specificity between Type I-F VchINT and Type V-K ShoINT systems, particularly in light of the significant off-target insertions previously observed for ShCAST^40^. After developing an alternative, unbiased NGS approach to query genome-wide integration events, which does not require the MmeI restriction enzyme used in Tn-seq, we first verified that this random fragmentation-based method returned similar specificity information for VchINT (**Supplementary Fig. 6a**). When the same method was applied to ShoINT and ShCAST, we found that only ~5–50% integration events were on-target, with substantial numbers of insertions distributed randomly across the genome (**Supplementary Fig. 6b,c**). These experiments highlight the remarkable fidelity of Type I-F INTEGRATE systems, and the need for further mechanistic studies to dissect the molecular basis of promiscuous integration by Cas12k-associated transposases. It will also be interesting to investigate the evolutionary forces that shaped I-F and V-K transposons, since one can envision competing selective pressures to spread widely without restrictive targeting constraints, while retaining enough specificity to mitigate fitness costs on the host.

### Single-step multiplex DNA insertions using INTEGRATE

CRISPR–Cas systems are naturally capable of multiplexing because of the way that CRISPR arrays are transcribed and processed, and transposase-mediated DNA integration exhibits intrinsic compatibility with multiple different genomic target sites as there is no requirement for target-specific homology arms. Thus, this provides a unique potential for multi-spacer CRISPR arrays to direct integration of the same cargo at multiple genomic targets simultaneously (**Fig. 4a**), which significantly reduces time and complexity for strain engineering projects requiring multi-copy integration. We explored this by first cloning a series of multiple-spacer arrays into pSPIN, and found that the integration efficiency of a *lacZ*-specific crRNA was unchanged for two spacers and reduced by <2-fold for three spacers, depending on relative position, when cells were cultured at 37 °C (**Fig. 4b**). Tn-seq analyses with double- and triple-spacer arrays revealed >99% on-target transposition, with characteristics that were otherwise indistinguishable from singleplex insertions for each target site (**Fig. 4c** and **Supplementary Fig. 7**), and we further verified multiplex insertions by whole-genome SMRT sequencing of double- and triple-insertion clones (**Supplementary Table 2**).

**Fig. 4.**
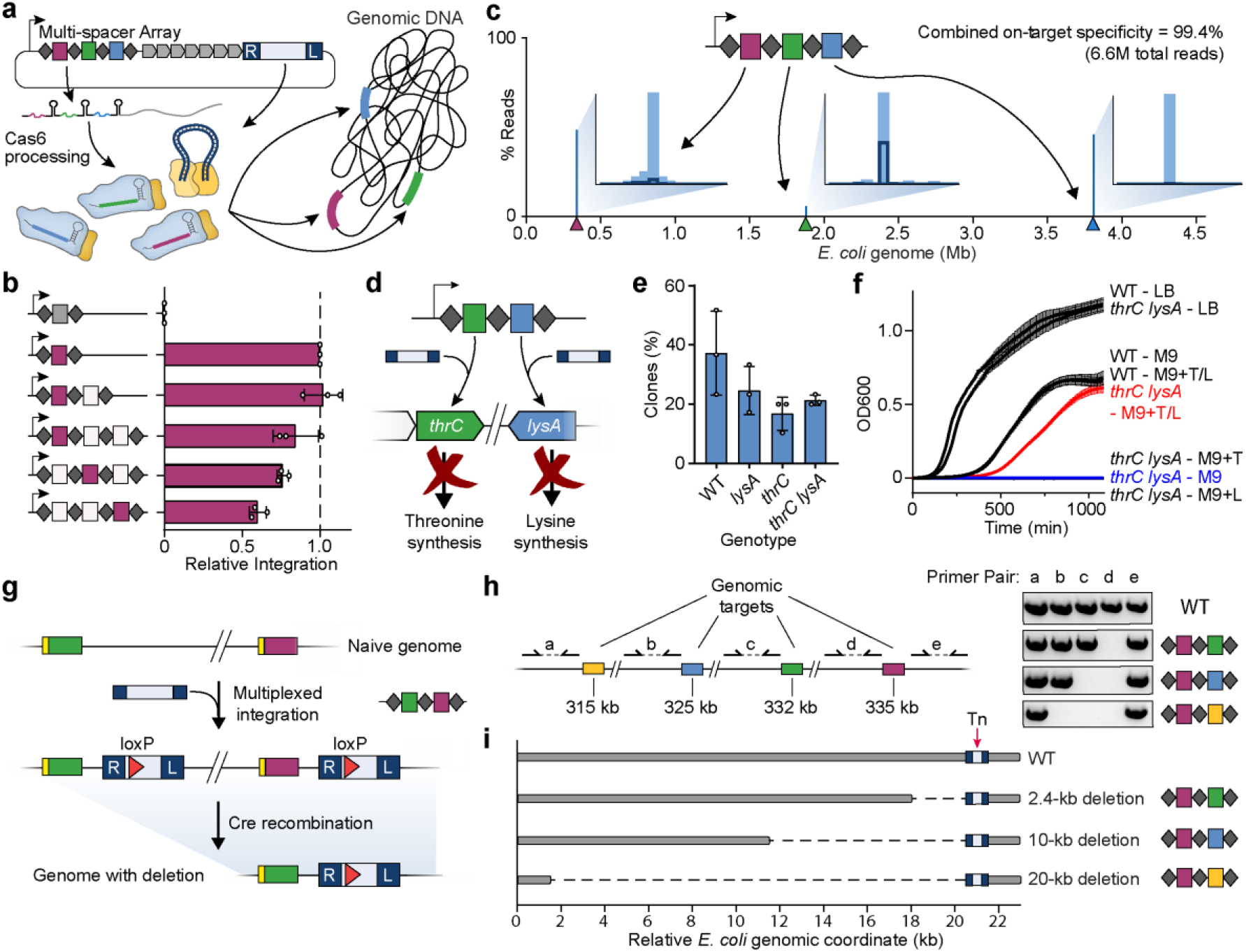
Multi-spacer CRISPR arrays direct multiplex insertions in a single step. **a,** Schematic of multiplexed RNA-guided DNA integration events with pSPIN encoding a multi-spacer CRISPR array. **b,** qPCR-based quantification of integration efficiency with crRNA-NT (grey) and crRNA-4 (maroon), encoded in a single-, double-, or triple-spacer CRISPR array in the position indicated; white squares represent other functional crRNAs. Data are normalized to the single-spacer array efficiency. **c,** Tn-seq data for a triple-spacer CRISPR array, plotted as the percent of total genome-mapping reads. The target sites are denoted by colored triangles, and the insets show the distribution of integration events within a 42–58 bp window downstream of the target site. **d,** Schematic of experiment using *thrC*- and *lysA*-specific spacers for single-step generation of threonine-lysine auxotrophic *E. coli*. **e,** Recovery percentage of the indicated clonal genotypes (WT, single-knockout, or double-knockout) after transforming *E. coli* with pSPIN encoding a double-spacer CRISPR array containing both *thrC*- and *lysA*-specific spacers. Colonies were stamped onto M9-agar supplemented with either H_2_O, threonine, lysine, or threonine and lysine, and genotypes were determined based on survivability in each of these conditions. **f,** Growth curves for WT and double-knockout *E. coli* clones cultured at 37 °C in LB or M9 minimal media with or without supplemented threonine (T) and lysine (L). **g,** Schematic of experimental approach to generate programmed genomic deletions. A double-spacer array directs multiplex insertion of two mini-Tn copies carrying LoxP sites; subsequent introduction of Cre recombinase leads to precise excision of the genomic fragment spanning the LoxP sites. **h,** Left, schematic showing genomic locus targeted for deletion. Right, the indicated double-spacer arrays were used to generate defined deletions, and PCR was performed with the indicated primer pairs to detect the presence/absence of genomic fragments flanking each target site, resolved by agarose gel electrophoresis. **i,** The programmed genomic deletions generated in **h** (2.4-, 10-, or 20-kb in length) were further verified by whole-genome, single-molecule real-time (SMRT) sequencing (**Methods**). The mini-transposon is indicated with a red arrow. Data in **b** and **e** are shown as mean ± s.d. for n = 3 biologically independent samples. Data in **f** are shown as mean ± s.d. for three technical replicates.

To further confirm that simultaneous insertions were indeed occurring within each individual chromosome rather than population-wide, we designed an experiment to generate auxotrophic *E. coli* strains requiring both threonine and lysine for viability by insertionally inactivating *thrC* and *lysA*^48^ (**Fig. 4d** and **Supplementary Fig. 8a–c**). Double-knockout clones could be rapidly isolated after a single transformation step (**Fig. 4e**), and exhibited selective growth in M9 minimal media only when both threonine and lysine were supplemented (**Fig. 4f**). To probe the stability of integration-based knockouts, we cultured clones in rich media for five serial overnight passages without removing the expression plasmid and observed no change in the media requirements (**Supplementary Fig. 8d**), thus confirming the potency of phenotypic outcomes driven by multiplexed INTEGRATE.

Finally, we explored the combined use of RNA-guided integrases with site-specific recombinases to mediate facile programmable, one-step genomic deletions. Specifically, we inserted a LoxP site within the mini-Tn cargo and generated double-spacer CRISPR arrays to drive multiplex integration at two target sites. Subsequently, we used Cre recombinase to excise the chromosomal region within the LoxP sites, thus resulting in a precise deletion containing a single mini-Tn (**Fig. 4g** and **Methods**). We designed CRISPR arrays to produce 2.4-, 10-, and 20-kb deletions, which we confirmed via diagnostic PCR analysis and unbiased, whole-genome SMRT sequencing (**Fig. 4h,i** and **Supplementary Fig. 9**). These experiments highlight the potential synergies of INTEGRATE with existing technologies, and the ease and versatility with which RNA-guided integrases can be leveraged for diverse and programmable genome-scale genetic modifications.

### Broad host-range activity of RNA-guided integrases

Mobile genetic elements, especially transposons, often ensure their evolutionary success by functioning robustly across a broad range of hosts, without a requirement for specific host factors^49^. Given this expectation, as well as the efficiency with which the *V. cholerae* machinery directs RNA-guided transposition in *E. coli*, we set out to evaluate INTEGRATE activity in other Gram-negative bacteria. We selected *Klebsiella oxytoca*, a clinically relevant pathogen implicated in drug-resistant infections^50^ and emerging model organism for biorefinery^51^, and *Pseudomonas putida*, an important bacterial platform for biotechnological and industrial applications^52,53^ (**Fig. 5a**). Using a pSPIN derivative driven by the constitutive J23119 promoter, we targeted four non-essential metabolic genes (*xylA, galK, lacZ,* and *malK*) and one antibiotic resistance gene (*ampR*) in *K. oxytoca*, as well as intergenic regions (upstream of *PP_2928* and *benR*) or genes previously edited (*nirC*, *nirD*, *bdhA*, and *PP_3889*) in *P. putida*^54,55^. For all 10 targets, we observed highly-accurate RNA-guided DNA integration by both PCR and Tn-seq, with similar integration distance and orientation bias profiles as seen in *E. coli* (**Fig. 5b,c** and **Supplementary Fig. 10a–d**). DNA insertions were virtually absent with a non-target crRNA, and on-target specificity was >95% on average, with the only outlier resulting from a prominent Cascade off-target binding site (**Supplementary Fig. 10e**). Given the potential for INTEGRATE to exhibit off-target activity similar to canonical CRISPR-Cas systems^56^, we developed a computational tool for guide RNA design and off-target prediction (**Supplementary Fig. 11)**; we anticipate updating this algorithm as additional mechanistic insights for programmable, RNA-guided integrases are acquired.

**Fig. 5.**
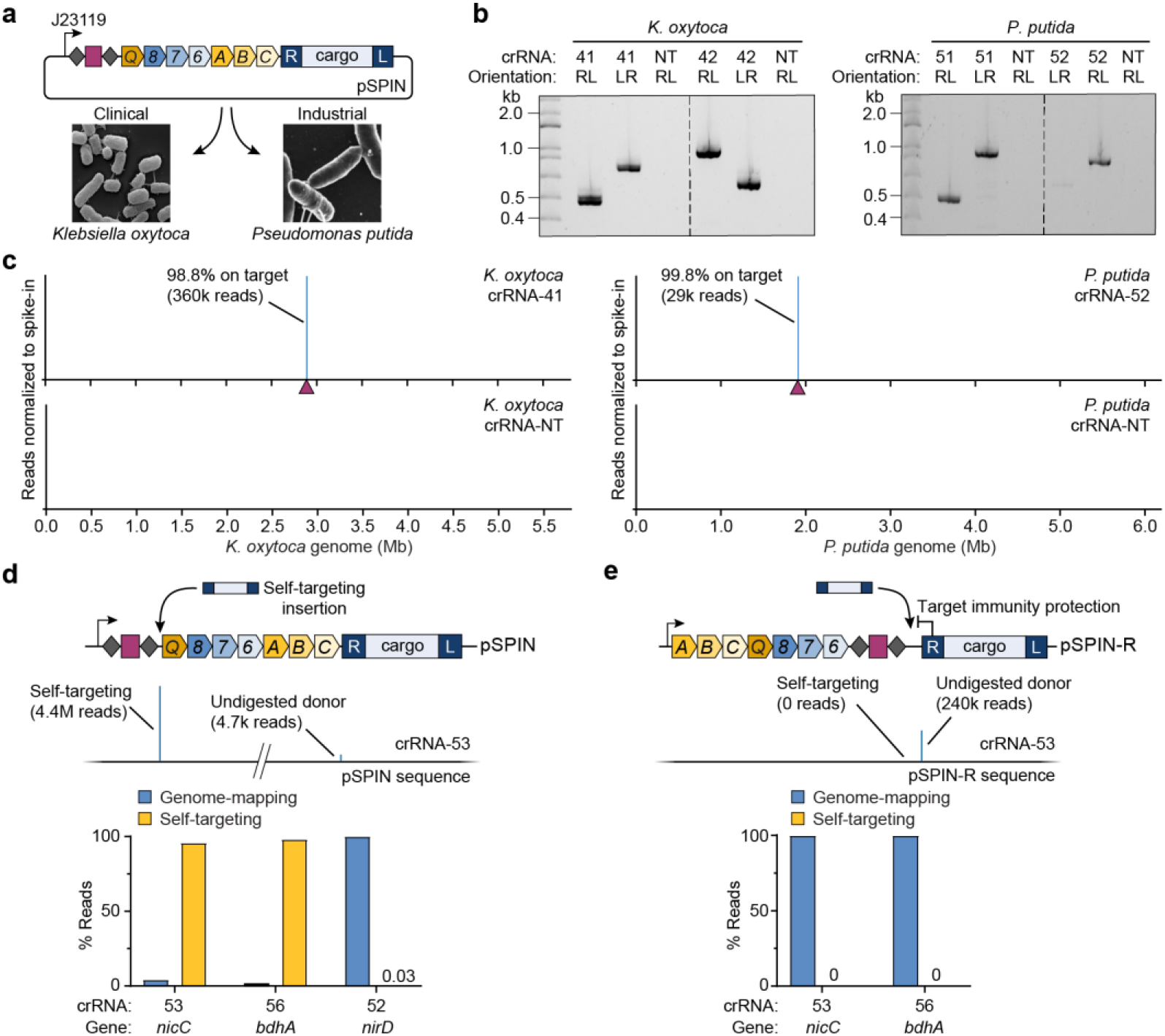
Robust and highly-accurate INTEGRATE activity in additional Gram-negative bacteria. **a,** Schematic showing the use of pSPIN constructs with constitutive J23119 promoter and broad-host pBBR1 backbone for RNA-guided DNA insertions in *Klebsiella oxytoca* and *Pseudomonas putida*. Micrographs taken from ^51,59^. **b,** PCR products probing for mini-Tn insertion at two different genomic loci in *K. oxytoca* (left) and *P. putida* (right), resolved by agarose gel electrophoresis. **c,** Normalized Tn-seq data for select targeting and non-targeting crRNAs for *K. oxytoca* (left) and *P. putida* (right). Genome-mapping reads are normalized to the reads from a spike-in control; the target site is denoted by a maroon triangle. **d,** Top, self-targeting of the spacer within the CRISPR array inactivates the pSPIN-encoded INTEGRATE system, and was detected for select crRNAs by Tn-seq (middle). Bottom, *P. putida* crRNAs targeting *nicC* and *bdhA*, but not *nirD*, show substantial plasmid self-targeting relative to genomic integration, as assessed by Tn-seq. **e,** Top, a modified vector (pSPIN-R) places the CRISPR array proximal to the mini-Tn, whereby self-targeting is blocked by target immunity. Bottom, *P. putida* crRNAs targeting *nicC* and *bdhA* no longer show any evidence of self-targeting with pSPIN-R, as assessed by Tn-seq.

Interestingly, for two of the *P. putida* crRNAs, we observed a substantial enrichment of NGS reads mapping to pSPIN, precisely 48-50 bp downstream of the spacer in the CRISPR array (**Fig. 5d**). We previously reported that Cascade-directed DNA integration exhibits a high degree of PAM promiscuity, including low activity with the mutant 5’-AC-3’ self-PAM present within CRISPR repeats flanking the spacer^35^. Indeed, we observed evidence of very low-level self-targeting in all of our *E. coli* Tn-seq datasets (**Supplementary Table 4**), and we suspected that the apparent abundance of self-targeting insertions for *P. putida* crRNAs targeting *nicC* and *bhdA* genes resulted from a fitness cost of the intended knockout and concomitant selective pressure to inactivate the pSPIN expression vector. We hypothesized that this ‘escape’ outcome could be avoided by redesigning the expression vector such that the self-target (i.e. the CRISPR array) would be in close proximity to the mini-Tn, and thus become protected by transposon target immunity. When we transformed *P. putida* with modified pSPIN-R vectors encoding the exact same crRNAs but at the 3’ end of the fusion transcript, we found that self-targeting was completely abrogated (**Fig. 5e** and **Supplementary Fig. 10f**).

Altogether, these experiments highlight the utility of RNA-guided integrases for programmable genetic modifications across diverse bacterial species, and illustrate how mechanistic knowledge of the transposition pathway can directly inform technology development efforts.

## Discussion

Through systematic engineering steps, we developed an optimized set of vectors to leverage INTEGRATE for targeted DNA integration applications in diverse bacterial species, without the need for DSBs, HR, or cargo-specific marker selection. These streamlined constructs can be easily modified to generate user-specific guide RNAs and genetic cargos, and they catalyze highly accurate, large DNA insertions at ~100% efficiency after a single transformation step. Moreover, by repurposing the natural CRISPR array, we demonstrate efficient multiplexing for simultaneous insertions or programmed genomic deletions by using INTEGRATE in combination with Cre-LoxP, within the same seamless workflow. Importantly, as target-specific homology arms are not required, the mini-Tn is compatible with any arbitrary target site, thus significantly reducing the complexity of the donor DNA and accelerating the cloning time, particularly for large-scale multiplex applications. This genetic engineering toolkit can be harnessed to generate large guide RNA libraries, which will enable high-throughput screening of rationally designed targeted DNA insertions that are not easily accessible with random transposase-based strategies. Furthermore, INTEGRATE can help advance existing strain engineering technologies, particularly those currently employing site-specific^23^ or non-specific^43,58^ transposases that could benefit from programmable site-specific insertions. Our observation of highly active integration at lower culturing temperatures provides a strategy for increasing the efficacy of genetic manipulations, and we anticipate that the broad temperature range of the system holds promise for general utility across diverse species. Finally, a deeper mechanistic understanding of transposition kinetics will also reveal how quickly integration is established within a bacterial cell population, potentially enabling future applications in microbial species that cannot be stably transformed with replicating plasmids.

Despite key advantages of RNA-guided DNA integration for bacterial engineering, we note some specific drawbacks that users should take into account. First, because transposon end sequences are essential for specific recognition by the transposase machinery, INTEGRATE is not suited for applications where precise, scarless insertions are required. Thus, generation of specific point mutations within endogenous genes, for example, will require gene editing- and recombination-based approaches, such as CRISPR-Cas9-coupled recombineering^8,14^. However, many other applications are not constrained by relatively short cargo-flanking sequences, such as simple insertional gene knockouts, or stable transgene integration into safe harbor regions. In addition, future transposon engineering may enable further reductions in the size of transposon ends, or their conversion into functional parts, such as peptide linkers for in-frame gene tagging. Second, applications involving iterative DNA insertions will need to carefully consider transposon target immunity and/or the risk of pre-existing insertions being mobilized by their cognate transposition machinery. While these effects may affect efficiencies, iterative insertions can still proceed using the same VchINT system, followed by validation of clones. However, orthogonal INTEGRATE systems provide avenues for circumventing these potential issues. The orthogonality of the *V. cholerae* and *S. hofmannii* INTEGRATE systems serves to illustrate the potential of multiple phylogenetically-distinct, cross-compatible transposases; as more CRISPR-transposon systems are discovered and functionally validated for both Type I and Type V, we envision the INTEGRATE toolkit expanding into a robust set of programmable, RNA-guided integrases that act orthogonally and are fully cross-compatible.

In addition to its utility for strain engineering, INTEGRATE systems may be a particularly useful tool in mixed bacterial communities and microbiome niches. Using our compact construct designs, we generated a fully autonomous CRISPR-transposon that was capable of high-efficiency integration, and in future studies, we envision mobilizing similar constructs on broad host-range conjugative plasmids, pre-programmed with multiple-spacer CRISPR arrays, to genetically modify desired bacterial species at user-defined target sites. Such self-driving systems would leverage the natural ability of transposons to propagate both within and between host genomes, while maintaining tight experimental control over specificity. This system will enable future gene drive applications, such as inactivating antibiotic resistance genes or virulence factors^57^ and introducing genetic circuits and synthetic pathways in a targeted manner.

## METHODS

### Plasmid construction

All *V. cholerae* INTEGRATE plasmid constructs were generated from pQCascade, pTnsABC, and pDonor, as described previously^35^, using a combination of restriction digestion, ligation, Gibson assembly, and inverted (around-the-horn) PCR. All PCR fragments for cloning were generated using Q5 DNA Polymerase (NEB).

Different plasmid backbone versions of pSPIN were cloned by generating one PCR fragment of the single INTEGRATE transcript and donor and combining with a digested vector backbone in a Gibson assembly reaction. pSPAIN was generated by Gibson assembly; a 0.98-kb mini-Tn was first inserted into a digested empty pBBR1 backbone, followed by double digestion of the cargo within the mini-Tn and insertion of the single INTEGRATE transcript.

ShoINT system was synthesized by GenScript; Cas12k and the sgRNA were synthesized as two separate cassettes on a pCDFDuet-1 (pCDF) plasmid, TnsA-TnsB-TniQ was synthesized as a native operon on a pCOLADuet-1 (pCOLA) plasmid, and the mini-Tn was synthesized on a pUC19 plasmid. Sho-pEffector and Sho-pSPIN were generated from these plasmids using Gibson assembly. ShCAST system was synthesized by GenScript according to the constructs described previously^40^, with pHelper on pUC19 and pDonor on pCDF backbones. Pairwise protein sequence similarities between the VchINT, ShoINT, and ShCAST machinery can be found in **Supplementary Table 3**.

Each construct containing a spacer was first constructed with a filler sequence containing tandem BsaI recognition sites in place of the spacer for VchINT and ShoINT, and tandem BbsI sites for ShCAST. New spacers were then cloned into the arrays by phosphorylation of oligo pairs with T4 PNK (NEB), hybridization of the oligo pair, and ligation into double BsaI- or BbsI-digested plasmid. Double- and triple-spacer arrays were cloned by combining two or three oligoduplexes with compatible sticky ends into the same ligation reaction. crRNAs for VchINT were designed with 32-nt spacers targeting sites with 5’ CC PAM. sgRNAs for ShoINT and ShCAST were designed with 23-nt spacers targeting sites with 5’ RGTN PAM and 5’ NGTT PAM, respectively. All spacer sequences used for this study are available in **Supplementary Table 5**. We note that our guide RNA design algorithm (**Supplementary Fig. 11**) was not used to generate spacers for this study.

Cloning reactions were transformed into NEB Turbo *E. coli*, and plasmids were extracted using Qiagen Miniprep columns and confirmed by Sanger sequencing (GENEWIZ). Transformed cells were cultured in liquid LB media or LB agar media, with addition of 100 μg/ml carbenicillin for pUC19 plasmids, 50 μg/ml spectinomycin for pCDF and pSC101*, and 50 μg/ml kanamycin for pCOLA, pSC101 and pBBR1. All plasmid construct sequences are available in **Supplementary Table 1**, and a subset are deposited at Addgene.

### *E. coli* culturing and general transposition assays

A full list of *E. coli* strains used for transposition experiments is provided in **Supplementary Table 6**.

All *E. coli* transformations were performed using homemade chemically competent cells and standard heat shock transformation, followed by recovery in LB at 37 °C and plating on LB-agar media with the appropriate antibiotics at the concentrations described above. Typical transformations efficiencies were >10^3^ CFU/μg of total DNA. All standard transposition assays in *E. coli* involved incubation at 37 °C for 24 hours after recovery and plating. However, experiments involving incubation at 30 °C or 25 °C were performed for an extended total of 30 hours; due to the reduced growth rate at these temperatures, cells were incubated longer to produce enough cell material for downstream analyses. To control for this extended incubation time, all incubations for Fig 4c were performed uniformly for 30 hours, including the 37 °C incubations. For similar reasons, incubation for the *ΔrecA* transposition assays (**Supplementary Fig. 4c**) was performed for 30 hours due to significantly slower growth rate of the *ΔrecA* strain.

For most experiments involving an IPTG-inducible T7 promoter, transformed cells were plated directly on 0.1 mM IPTG LB agar plates for 24 hours after recovery. Exceptions were for the pUC19 pSPIN construct (Fig 1d), all transposition assays performed for Fig 1e, and all ShoINT and ShCAST experiments - where transformed cells were first plated on LB agar without induction and incubated for 16 hours, and were then scraped and replated on LB agar with 0.1 mM IPTG and incubated for 24 hours. For these experiments, this replating protocol was generally used since initial transformation efficiencies were low, potentially from significant IPTG induced toxicities affecting transformation; separating transformation and induction steps allowed for enough cells to be generated for lysis and further analyses. To avoid IPTG degradation affecting transposition efficiency, LB agar plates were made fresh with frozen IPTG stocks, and were kept at 4 °C and used within 7 days of preparation. All post-transformation incubations involving active transposition were performed on solid media to avoid competitive growth effects causing enrichment of rare events.

Experiments involving three plasmids - pDonor, pTnsABC and pQCascade or variants - were performed by first transforming pTnsABC and pDonor into chemical competent cells, picking a single colony and growing overnight in liquid LB media with double antibiotic selection, inducing chemical competency using standard methods, and then transforming these cells with the pQCascade plasmid. Experiments involving two plasmids were performed by co-transformation of both plasmids into chemical competent cells simultaneously, although we note that this generally resulted in lower transformation efficiencies and required more input DNA than if the plasmids had been transformed iteratively.

### Transposition assays in *Klebsiella oxytoca and Pseudomonas putida*

A full list of bacterial strains used for transposition experiments is provided in **Supplementary Table 6**.

For *K. oxytoca* transformations, cells were grown overnight to saturation, and were diluted 1:100 and grown to OD600 of ~0.4-0.5. Cells were then placed on ice for 15-30 min and subsequently washed three times with ice-cold 10% glycerol DI water. After the washes, cells were concentrated 100-fold in ice-cold 10% glycerol DI water. 50 μl of cells were electroporated with 50 ng plasmid, using 0.1 cm cuvettes at 1.8 kV. Cells were recovered in 1 ml of LB media for 2 hours at 37 °C, and were plated on LB agar with selection at 37 °C for 24 hours.

For *P. putida* transformations, we adapted a previously described protocol^58^. Briefly, overnight cultures were washed three times with 300 mM sucrose and concentrated 50-fold. Cells were then distributed into 100 μl aliquots and separately electroporated with 100 ng of plasmid using 0.2 cm cuvette at 2.5 kV, and were recovered in 1 ml of LB media for 2 hours at 30 °C. Recovered cells were plated on LB agar with selection at 30 °C for 24 hours.

All transposition assays for *K. oxytoca* and *P. putida* were performed by transforming a pSPIN construct on a pBBR1 backbone, expressed from a constitutive J23119 promoter. Cells were incubated on LB agar for 24 hours after recovery; colonies were then scraped for gDNA extraction using the Wizard Genomic DNA Purification kit (Promega).

### PCR and qPCR analysis of transposition

*E. coli* cells transformed with INTEGRATE machinery were scraped from LB agar plates and suspended in liquid LB, and the OD600 of the resulting suspensions were taken. From each resuspension, approximately 3.2 × 10^8^ cells (equivalent to 200 μl of OD600 = 2.0 of resuspended cells) were taken for lysis. In scenarios where colonies were small and less than this amount of cell resuspension was recovered, the entire resuspension of cells was used for lysis. Cells were pelleted by centrifugation at 4000*g* for 2 min, the LB supernatant was poured off and cells were resuspended in 80 μl of DI water, followed by lysis at 95 °C for 10 min. The lysates were cooled to room temperature, pelleted by centrifugation at 4000*g* for 2 min, and the supernatant was diluted 20-fold in DI water and used for subsequent analyses. Further dilutions of lysates may be used for analysis, while we have observed polymerase inhibition from raw lysates at higher concentrations than the 20-fold dilution, especially for qPCR.

PCR reactions for *E. coli* samples were performed using Q5 Polymerase (NEB) in a 12.5 μl reaction containing 200 μM dNTPs, 0.5 μM of each primer, and 5 μl of diluted lysate supernatant. Primer pairs involved one mini-Tn-specific primer and one genome-specific primer, and each primer pair probes for integration in either T-RL of T-LR orientation. PCR amplicons were generated over 30 PCR cycles, and were resolved by gel electrophoresis on 1-1.5% agarose stained with SYBR Safe (Thermo Scientific). PCR reactions for *K. oxytoca* and *P. putida* were done using similar primer design as E. coli, with Q5 Polymerase in a standard 50 μl reaction mixture, and with 20 ng extracted gDNA as input instead of cell lysate.

qPCR reactions were performed on 2 μl of diluted lysates in 10 μl reactions, containing 5 μl SsoAdvanced Universal SYBR Green 2X Supermix (BioRad), 2 μl of 2.5 μM mixed primer pair, and 1 μl H_2_O. Each lysate sample was analyzed with 3 separate qPCR reactions involving 3 primer pairs, as described previously^35^: two pairs each involving one mini-Tn-specific primer with one genomic-specific primer probing for either the T-RL or T-LR integration orientation, and one pair with two genome-specific reference primers at the rssA locus. Primer pairs were designed to amplify a product between 100 – 250 bp, and were confirmed to have amplification efficiencies between 90%-110% using serially diluted lysates. A full list of qPCR primers used in this study is provided in **Supplementary Table 7**. Integration efficiency (%) for each insertion orientation is defined as 100 × (2^ΔCq), where ΔCq is the Cq(genomic reference pair) - Cq(T-RL pair OR T-LR pair); total integration percentage is the sum of both orientation efficiencies.

We note that our qPCR protocol has previously been benchmarked using lysate samples simulating known integration efficiencies and orientation biases^35^. However, as efficiency is dependent on Cq measurement of both the genomic reference primer pair and the integration junction primer pair, variation in the measurement of either impacts the final efficiency value. This can lead to apparent measurements of >100% when the actual integration efficiency is close to 100%, particularly since variations in raw Cq values are amplified as the magnitude of ΔCq increases. Thus, qPCR noise affects efficiency measurements more disproportionately at higher efficiencies.

### Isolation of clonally-integrated *E. coli* colonies

As we previously described the potential for colonies becoming polyclonal as integration occurs alongside colony expansion^35^, all clonal isolation steps were preceded by a “bottlenecking” step, where all colonies were scraped, resuspended in LB, and plated at an appropriate dilution to obtain a new set of colonies. Colonies were then picked and resuspended in 100 μl of MQ water, followed by lysis at 95 °C for 10 min. 5 μl of lysate was then used as input template for Q5 PCR as described above. Colonies were identified as clonal using three sets of PCRs per target site per lysate, as described previously^35^. Briefly, two PCR pairs probed for the presence of either T-RL or T-LR integration, respectively, and a third pair amplified across the genomic region of the expected insertion junction. A colony was considered clonal when only one of the first two primer pairs leads to amplification, and the third pair amplified solely a larger product that corresponds to the genomic region plus the mini-Tn. Where crRNA-4 (targeting *lacZ*) was used for integration, blue-white screening was used to select for white colonies, which were then confirmed with the above PCR strategy.

### VchINT target immunity experiments

A pSPIN derivative with crRNA-4 (targeting *lacZ*) on a pSC101* temperature-sensitive backbone was used to insert a 0.98 kb mini-Tn into BL21(DE3) cells at 30 °C for 30 hours. A clonal-insertion strain was isolated as described above, and the pSPIN plasmid was cured by culturing cells at 37 °C overnight in liquid LB media. Resulting cells were made chemically competent, and a separate pDonor containing a different cargo was transformed alongside a pEffector construct with a crRNA targeting a site *d* (bp) away from the original crRNA-4 target (as indicated in **Fig. 3a**). qPCR was then performed, where Tn-specific primers were designed to bind in the cargo in order to distinguish it from the original crRNA-4 insertion. For each target site, normalization was done by performing the same transposition and qPCR assay in WT BL21(DE3) cells, and dividing the immunized qPCR efficiency by the WT efficiency. We note that, due to the presence of two identical repeats of the mini-Tn right end and left end (111 bp and 149 bp, respectively) from the original and new insertions, it is possible that the observed target immunity phenotype is affected by low-level recombination between these repetitive sequences, which is not taken into account in our analyses.

### Mini-Tn remobilization experiments

BL21(DE3) cells with a clonal crRNA-4 (*lacZ*) insertion, isolated and cured of INTEGRATE plasmids as described above, was made chemically competent. A pEffector construct with crRNA-1 (targeting downstream of *glmS*) was transformed into these cells, without a donor plasmid containing a new mini-Tn. Presence of the mini-Tn at both *lacZ* and *glmS* was probed for by PCR as described above.

Mini-Tn-competition experiments were set up similarly, where a pEffector construct with crRNA-1 was transformed along with a pDonor which carries the same mini-Tn as the *lacZ*-insertion, except for a 5-bp mutation at the 3’ end of the R-end. This mutation site was used to design Tn-specific primers to distinguish the genomic-insertion and plasmid-borne mini-Tn at both *lacZ* and *glmS* sites.

### VchINT/ShoINT orthogonality experiments

For the orthogonality experiments in **Fig. 3c**, BL21(DE3) cells were co-transformed with a two-plasmid combination of either Vch-pEffector or Sho-pEffector, and either Vch-pDonor or Sho-pDonor. The spacers for both systems were designed to target the same region of the *lacZ* locus. For PCR analysis of integration activity, due to the two pDonors carrying the same cargo, transposon-specific primers were designed to bind in the R-end or L-end of the mini-Tn.

For data shown in **Fig. 3d**, Sho-pEffector and Sho-pDonor were co-transformed into BL21(DE3) containing a clonal *lacZ*-insertion. The Sho-sgRNA was designed with a spacer targeting a similar region near the *glmS* locus that is targeted by Vch-crRNA-1. PCR analysis was performed as described above.

### Amino acid auxotrophy experiments

M9 minimal media was prepared with the following components: 1X M9 salts (Difco), 0.4% glucose, 2 mM MgSO_4_, and 0.1 mM CaCl_2_. M9 agar was prepared as above, with the addition of 15 g/l of Dehydrated agar (BD). L-threonine and/or L-lysine was supplemented at 1 mM as indicated.

For individual *thrC* or *lysA* targeting experiments, BL21(DE3) cells were transformed with a pSPIN construct with a crRNA targeting either gene. Transformed cells were incubated on LB agar at 37 °C for 24 hours. Bottlenecking and clonal insertions identification by PCR were performed as described above, and cells were then evaluated for ability to grow in M9 minimal media with and without addition of the appropriate amino acid.

For multiplexed targeting of both *thrC* and *lysA*, BL21(DE3) cell were transformed with a pSPIN construct expressing a *thrC*-*lysA*-targeting double-spacer array. Cells were then incubated and bottlenecked on LB agar as above, and bottlenecked colonies were then stamped onto M9 agar plates supplemented with either no amino acids, only threonine or lysine, or both amino acids, to identify growth phenotype. For data presented in **Fig. 4e**, this screen was performed on 30 colonies for each of three independent experiments.

OD600 growth curve analysis was performed by first inoculating WT BL21(DE3) or isolated auxotrophic strains from −80 °C glycerol stocks into LB media for overnight growth. 1 ml of each culture was then pelleted at 16000*g* and resuspended in 1 ml MQ water, and was inoculated at a 1:1000 dilution into the respective growth media on a 96-well cell culture plate. Growth assay was then performed with a Synergy H1 plate reader shaking at 37 °C for 18 hours, and OD at 600nm taken every 5 min. Each sample was measured in three technical-replicates in separate wells on the sample plate, and were normalized to blank wells containing media only.

### Cre-Lox genomic deletion experiments

BL21(DE3) cells were transformed with a pSPIN construct containing a double-spacer CRISPR array containing crRNA-4 and a second spacer targeting the same strand either 2.4-, 10- or 20-kb away from crRNA-4. The mini-Tn of this construct was previously modified to include a 34-bp recognition sequence for Cre recombinase. Cells were incubated and bottlenecked, and colonies with double-clonal insertions were isolated by a combination of blue-white screening and PCR, as described above. We note that, although the two targets for the 2.4-kb deletion were within each other’s range for target immunity effects, we were still readily able to isolate the desired clones. Double-insertion clones were made chemically competent, and were then transformed with a plasmid expressing Cre recombinase from an IPTG-inducible T7 promoter. Cells were incubated at 37 °C for 16 hours and bottlenecked, and colonies having undergone recombination were isolated by PCR. We observed small colonies and very low transformation efficiencies when transformed cells were plated on 0.1 mM IPTG, while we were readily able to isolate recombined clones without IPTG induction, suggesting that small amounts of Cre resulting from leaky T7 expression were sufficient for recombination. Thus, all Cre-recombinase transformations were performed with no IPTG present.

### Tn-seq library preparation and sequencing

Transformations for Tn-seq transposition assays were carried out as described above, using donor plasmids containing a mini-Tn where the 8-nt terminal repeat of the mini-Tn R-end was mutated to contain an MmeI recognition sequence (from 5’-TGGTGATA to 5’-TGGTG<u>GA</u>A). We previously showed that a mini-Tn with this mutation is still functionally active, with a ~50% decrease in total integration efficiency^35^. Transformed cells were incubated on LB agar at 37 °C for 24 hours, except for assays shown in **Supplementary Fig 4d,f**, where cells were incubated at 30 °C for 30 hr. Colonies were then scraped and resuspended in liquid LB media, and 0.5 ml (approximately 2 × 10^9^ cells) were used for gDNA extraction with the Wizard Genomic DNA Purification kit (Promega), which typically yielded 50 μl of 0.5-1.5 μg/μl gDNA.

NGS libraries were prepared in parallel in PCR tubes, each with 1 μg of gDNA first being digested with 4 U of MmeI (NEB) for 2 hours at 37 °C, in a 50 μl reaction containing 50 μM S-adenosyl methionine and 1x CutSmart buffer, followed by heat inactivation at 65 °C for 20 min. MmeI digestion results in the generation of 2-nucleotide 3’-overhangs. Reactions were cleaned up with 1.4X Mag-Bind TotalPure NGS magnetic beads (Omega) according to the manufacturer’s instructions, and elutions were done using 30 μl of 10 mM Tris-Cl, pH 7.0. Double-stranded i5 universal adaptors containing a 3’-terminal NN overhang were ligated to the MmeI-digested gDNA in a 20 μl ligation reaction consisting of 16.86 μl of MmeI-digested gDNA, 5 nM adaptor, 400 U T4 DNA ligase (NEB), and 1X T4 DNA ligase buffer. Reactions were left at room temperature for 30 min, and were then cleaned with magnetic beads. Since the donor plasmid (either pDonor, pSPIN or pSPIN-R) contains a copy of the mini-Tn that can also be digested with MmeI and ligated with i5 adaptor, we included a restriction enzyme recognition site (HindIII for pDonor, or Bsu36I for pSPIN and pSPIN-R) in the 17-bp space between the 5’ end of the mini-Tn and the MmeI digestion site. This allowed us to reduce contamination of donor sequences within the NGS libraries, by digesting the entirety of the adaptor-ligated gDNA elution with 20 Units of HindIII or Bsu36I in a 34.4 μl reaction for 2 hours at 37 °C, before a heat inactivation step at 65 °C for 20 min. DNA clean-up using magnetic beads was then performed.

Eluted DNA was then amplified in a PCR-1 step, where adaptor-ligated transposons were enriched using a universal i5-adaptor primer and a transposon-specific primer with a 5’ overhang containing a universal i7 adaptor. In a 25 μl PCR-1 reaction, 16.7 μl of HindIII/Bsu36I-digested gDNA was mixed with 200 μM dNTPs, 0.5 μM primers, 1X Q5 reaction buffer, and 0.5 U Q5 DNA Polymerase (NEB). Amplification proceeded for 25 cycles at an annealing temperature of 66 °C. 20-fold dilutions of the reaction products were used as template for a second 20 μl PCR reaction (PCR-2) with indexed p5/p7 Illumina primers. The PCR-2 reaction was subjected to 10 additional amplification cycles with an annealing temperature of 65 °C, after which analytical gel electrophoresis was performed to verify amplification for each library. Barcoded reactions were pooled and resolved by 2.5% agarose gel electrophoresis, followed by isolation of DNA using Gel Extraction Kit (Qiagen), and NGS libraries were quantified by qPCR using the NEBNext Library Quant Kit (NEB). Illumina sequencing was performed with a NextSeq mid-output kit with 150-cycle single-end reads and automated adaptor trimming and demultiplexing (Illumina).

For pSPIN libraries involving a spike-in, 10 μl of a 0.02 ng/μl spike-in plasmid was added to each 1 μg DNA sample prior to MmeI digestion, and library preparation proceeded as described above. The plasmid contains a full-size MmeI-mini-Tn, where there is no Bsu36I restriction site in the 17-bp fingerprint space – thus this fingerprint survives the Bsu36I donor digestion step for pSPIN libraries, and provides a constant “contamination” into the library to control for sequencing depth.

### Random fragmentation library prep and sequencing

BL21(DE3) cells were transformed with Vch-pSPIN or Sho-pSPIN, or were co-transformed with pHelper and pDonor for ShCAST. Transformation, incubation and gDNA extraction with the Wizard Genomic DNA Purification kit (Promega) were performed as described previously.

Following the NEBNext® dsDNA Fragmentase protocol, about 2.5 μg of gDNA was fragmented for 14 min. The fragmentation reactions were purified using 1.4X Mag-Bind® Total Pure NGS (Omega) beads with an elution step in 30 μl 1X TE. Approximately 1 μg of the fragmented DNA was used for end preparation, adapter ligation and USER cleavage, according to the NEBNext Ultra II DNA Library Prep Kit for Illumina protocol. The reactions were purified using 1.2X Mag-Bind® Total Pure NGS (Omega) beads with an elution step in 30 μl MQ water.

To reduce the number of fragments deriving from the mini-Tn on the donor plasmid, the samples were digested with restriction enzymes (VchINT - KpnI/Bsu36I, ShoINT - PstI/HindIII, ShCAST - NcoI/AvrII) overnight at 37 °C. The reactions were then purified using 1.2X Mag-Bind® Total Pure NGS (Omega) beads with an elution step in 30 μl MQ water.

PCR-1 reactions were performed using Q5 Polymerase (NEB) in a 20 μl reaction containing 200 μM dNTPs, 0.5 μM of each primer, and 30 ng of input DNA. A transposon-specific primer carrying an i5 adapter, and an i7-specific primer specifically amplified transposon containing fragments over 20 PCR cycles. A second PCR reaction (PCR-2) was used to add specific Illumina index sequences to the i5 and i7 adapters over 10 PCR cycles in a 25 μl reaction with 1.25 μl from PCR-1 as the input DNA.

Samples were purified using the Qiagen PCR Clean-up Kit, and their DNA concentrations were measured using a DeNovix spectrometer. The amount of DNA was normalized and samples were combined. The pooled libraries were then quantified using the NEBNext® Library Quant Kit for Illumina, and Illumina sequencing was performed as described above.

### Analysis of NGS data

All analyses of Tn-seq and random fragmentation sequencing data were performed using a custom Python pipeline. Demultiplexed raw reads were filtered to remove reads where less than half of the bases passed a Phred quality score of 20 (Q20 – corresponding to >1% base miscalling). Reads that contained the 15-bp 5’ terminal sequence of the mini-Tn R-end (allowing up to one mismatch) were then selected, and the 17-bp sequence directly upstream of this R-end sequence was extracted. This 17-bp “fingerprint” sequence corresponds to the distance from the R-end to the MmeI digestion site, and contains the sequence context in which the mini-Tn is found (**Supplementary Fig. 3a**). Reads without sufficient length to extract a 17-bp fingerprint were removed from analysis. For each random fragmentation sample, since the two transposon ends were amplified and sequenced as two separate libraries, extraction of fingerprints from reads were performed separately for the R and L transposon ends.

Fingerprint sequences were aligned to reference genomes of the corresponding species and strain, depending on each specific library. The full list of strains, species, and corresponding reference genome accession identifiers is provided in **Supplementary Table 6**; reference genomes for *E. coli* and *P. putida* were obtained from published NCBI genomes, whereas our *K. oxytoca* parent strain was sequenced and assembled de novo using whole-genome SMRT sequencing to obtain the reference genome (see below for SMRT sequencing method). Alignment to reference was performed using the bowtie2 alignment library - perfect mapping was used for alignment, and only reads that aligned exactly once to the reference genome were used for downstream analyses. Fingerprints that did not map to the reference genome were screened for sequences corresponding to undigested donor contamination, or for fingerprints mapping downstream of the CRISPR array on the donor plasmid, which correspond to self-targeting events (**Fig. 5d,e**). For cases where a spike-in plasmid was used, the number of fingerprints containing the spike-in sequence was also determined.

Bowtie2 alignment outputs were used to generate genome-wide integration distributions, the number of reads corresponding to integration events at each position across the reference genome was plotted. For visualization purposes, these positions were grouped into 456 separate 10-kb bins, and peaks were plotted as a percentage of total reads. In cases where a spike-in was used, peaks were further normalized by the number of spike-in fingerprints detected, and the plot each non-targeting control was plotted to the same y-axis scale as its corresponding targeting sample. This analysis was performed similarly for each random fragmentation library by combining R-end and L-end fingerprints prior to alignment and plotting.

Integration-site distance distribution plots were generated from bowtie2 alignments by plotting number of reads against the distance between the 3’ end of the protospacer and the site of insertion corresponding to the reads, at single-bp resolution. The on-target % was calculated as the percentage of reads corresponding to integration events within a 100-bp window centered at the integration site with the largest number of reads. The orientation bias of integration which we define as the ratio of number of reads corresponding to T-RL insertions to those corresponding to T-LR insertions. For random fragmentation libraries, alignments for this analysis were performed separately for R-end and L-end fingerprints, and the results were combined to generate the plot.

We note that our Tn-seq sequencing is susceptible to potential biases arising from differences in MmeI digestion efficiency at each site, and in ligation efficiencies of 3’-terminal NN overhang adaptors, which were not taken into account by our downstream analyses.

### PacBio SMRT sequencing and analysis

gDNA samples for library preparation were extracted from overnight LB cultures using the Wizard Genomic DNA Purification kit (Promega) as described above. Multiplexed microbial whole genome SMRTbell libraries were prepared as recommended by the manufacturer (Pacific Biosciences). Briefly, two micrograms of high molecular weight genomic DNA from each sample (n=12 per pool) was sheared using a g0-tube to ~10kb (Covaris). These sheared gDNA samples were then used as input for SMRTbell preparation using the Template Preparation Kit 1.0, where each sample was treated with a DNA Damage Repair and End Repair mix, in order to repair nicked DNA and repair blunt ends. Barcoded SMRTbell adapters were ligated onto each sample in order to complete SMRTbell library construction, and then these libraries were pooled equimolarly, with a final multiplex of 12 samples per pool. The pooled libraries were then treated with exonuclease III and VII to remove any unligated gDNA, and cleaned with 0.45X AMPure PB beads to remove small fragments and excess reagents (Pacific Biosciences). The completed 12-plex pool was annealed to sequencing primer V3 and bound to sequencing polymerase 2.0 before being sequenced using one SMRTcell 8M on the Sequel 2 system with a 20-hour movie.

After data collection, the raw sequencing reads were demultiplexed according to their corresponding barcodes using the Demultiplex Barcodes tool found within the SMRTLink analysis suite, version 8.0. Demultiplexed subreads were downsampled 10-fold by random downsampling, and assembled de novo using the Hierarchical Genome Assembly Process (HGAP) tool, version 4.0 using the following parameters: Aggressive mode = off, Downsampling factor = 0, Minimum mapped length = 50 bp, Seed coverage = 30, Consensus algorithm = best, Seed length cutoff = −1, Minimum Mapped Concordance = 70%.

Subread mapping and structural variant analysis were performed using the PB-SV tool within SMRTLink 8.0, using the BL21(DE3) genome (Accession CP001509.3) as reference, with the following parameters: Minimum SV length = 20 bp, Minimum reads supporting variant for any one sample = 2, Minimum mapped length = 50 bp, Minimum length of copy number variant = 1000 bp, Minimum reads supporting variant (total over all samples) = 2, Minimum % of reads supporting variant for any one sample = 20%, Minimum mapped concordance = 70%. VCF outputs was used to generate SV analysis results shown in **Supplementary Table 2**, and BAM alignments were visualized with IGV to generate genome-deletion coverage plots (**Supplementary Fig. 9**).

For the coverage plot of the 10-kb insertion (**Supplementary Fig. 4g**), circular consensus sequence (CCS) reads were generated with SMRTLink 8.0, and were then filtered using a custom Python script to obtain only reads containing 20 bp of the R-end and/or the L-end of the mini-Tn, where the 20-bp regions directly adjacent to these R/L-end sequences do not map to pDonor. These filtered reads were then aligned to an artificial reference genome, where the entire 10-kb mini-Tn was inserted 49 bp downstream of the crRNA-4 target sequence of the CP001509.3 reference genome, using Geneious Prime at medium sensitivity with no fine-tuning.

## Data availability

Next-generation sequencing data are available in the National Center for Biotechnology Information Sequence Read Archive (BioProject Accession:).

## Code availability

Custom Python scripts used for the described NGS data analyses are available online via GitHub (https://github.com/sternberglab/Illumina-pipeline/tree/vo2020Paper). The INTEGRATE guide RNA design tool and associated documentation are available online via GitHub (https://github.com/sternberglab/INTEGRATE-guide-RNA-tool)

## ACKNOWLEDGMENTS

We thank N. Jaber for laboratory support, J. Bondy-Denomy for discussions, L.F. Landweber for qPCR instrument access, J. Mohabir for assistance with the NGS read alignment, the JP Sulzberger Columbia Genome Center for NGS support, and M.L. Smith, I. Oussenko, and the Genomics Technology Laboratory at the Icahn School of Medicine at Mount Sinai for SMRT sequencing. H.H.W. acknowledges funding support from NSF (MCB-1453219), NIH (1U01GM110714, 1R01AI132403), ONR (N00014-17-1-2353), and the Burroughs Wellcome Fund (PATH1016691) for this work. C. R. is supported by a Junior Fellows Scholarship from the Simons Society of Fellows. S.H.S. acknowledges a generous start-up package from the Columbia University Irving Medical Center Dean’s Office and the Vagelos Precision Medicine Fund.

## AUTHOR CONTRIBUTIONS

P.L.H.V. and S.H.S. conceived of and designed the project. P.L.H.V. performed most experiments and analyzed the data. C.R. performed experiments in *K. oxytoca* and *P. putida*, with guidance from H.H.W. S.E.K. performed target immunity, ShoINT, and random fragmentation NGS experiments. E.E.C. helped with cloning and transposition experiments. C.A. assisted with computational analyses of NGS data and the guide RNA design algorithm. P.L.H.V., S.H.S., and all other authors discussed the data and wrote the manuscript.

## COMPETING INTERESTS

P.L.H.V., S.E.K., and S.H.S. are inventors on patents and patent applications related to CRISPR– Cas systems and uses thereof. H.H.W is a scientific advisor to SNIPR Biome. S.H.S. is a co-founder and scientific advisor to Dahlia Biosciences, and an equity holder in Dahlia Biosciences and Caribou Biosciences.

Correspondence and requests for materials should be addressed to S.H.S.

## REFERENCES

1. Dunbar, C. E. et al. Gene therapy comes of age. Science 359, eaan4672 (2018).

2. Gelvin, S. B. Integration of Agrobacterium T-DNA into the Plant Genome. Annu Rev Genet 51, 195–217 (2017).

3. Davy, A. M., Kildegaard, H. F. & Andersen, M. R. Cell Factory Engineering. Cell Systems 4, 262–275 (2017).

4. Brophy, J. A. N. et al. Engineered integrative and conjugative elements for efficient and inducible DNA transfer to undomesticated bacteria. Nature Microbiology 3, 1043–1053 (2018).

5. Miyazaki, R. & van der Meer, J. R. A new large-DNA-fragment delivery system based on integrase activity from an integrative and conjugative element. Appl Environ Microbiol 79, 4440–4447 (2013).

6. Martínez-García, E. & de Lorenzo, V. Transposon-based and plasmid-based genetic tools for editing genomes of gram-negative bacteria. Methods Mol Biol 813, 267–283 (2012).

7. van Opijnen, T., Bodi, K. L. & Camilli, A. Tn-seq: high-throughput parallel sequencing for fitness and genetic interaction studies in microorganisms. Nat Meth 6, 767–772 (2009).

8. Wang, H. H. et al. Genome-scale promoter engineering by coselection MAGE. Nat Meth 9, 591–593 (2012).

9. Sharan, S. K., Thomason, L. C., Kuznetsov, S. G. & Court, D. L. Recombineering: a homologous recombination-based method of genetic engineering. Nat Protoc 4, 206–223 (2009).

10. Zhang, Y., Buchholz, F., Muyrers, J. P. & Stewart, A. F. A new logic for DNA engineering using recombination in Escherichia coli. Nat Genet 20, 123–128 (1998).

11. Baba, T. et al. Construction of Escherichia coli K-12 in-frame, single-gene knockout mutants: the Keio collection. Mol Syst Biol 2, 2006.0008 (2006).

12. Cotta-de-Almeida, V., Schonhoff, S., Shibata, T., Leiter, A. & Snapper, S. B. A new method for rapidly generating gene-targeting vectors by engineering BACs through homologous recombination in bacteria. Genome Res 13, 2190–2194 (2003).

13. Datsenko, K. A. & Wanner, B. L. One-step inactivation of chromosomal genes in Escherichia coli K-12 using PCR products. Proc Natl Acad Sci USA 97, 6640–6645 (2000).

14. Jiang, W., Bikard, D., Cox, D., Zhang, F. & Marraffini, L. A. RNA-guided editing of bacterial genomes using CRISPR-Cas systems. Nat Biotechnol 31, 233–239 (2013).

15. Wang, K. et al. Defining synonymous codon compression schemes by genome recoding. Nature 539, 59–64 (2016).

16. Sukhija, K. et al. Developing an extended genomic engineering approach based on recombineering to knock-in heterologous genes to Escherichia coli genome. Mol. Biotechnol. 51, 109–118 (2012).

17. Wang, H. H. et al. Programming cells by multiplex genome engineering and accelerated evolution. Nature 460, 894–898 (2009).

18. Vento, J. M., Crook, N. & Beisel, C. L. Barriers to genome editing with CRISPR in bacteria. J. Ind. Microbiol. Biotechnol. 46, 1327–1341 (2019).

19. Jiang, Y. et al. CRISPR-Cpf1 assisted genome editing of Corynebacterium glutamicum. Nat Commun 8, 15179 (2017).

20. Kosicki, M., Tomberg, K. & Bradley, A. Repair of double-strand breaks induced by CRISPR-Cas9 leads to large deletions and complex rearrangements. Nat Biotechnol 36, 765–771 (2018).

21. Wannier, T. M. et al. Improved bacterial recombineering by parallelized protein discovery. bioRxiv 14, 1–69 (2020).

22. Corts, A. D., Thomason, L. C., Gill, R. T. & Gralnick, J. A. A new recombineering system for precise genome-editing in Shewanella oneidensis strain MR-1 using single-stranded oligonucleotides. Sci Rep 9, 39–10 (2019).

23. Peters, J. M. et al. Enabling genetic analysis of diverse bacteria with Mobile-CRISPRi. Nature Microbiology 4, 244–250 (2019).

24. St-Pierre, F. et al. One-step cloning and chromosomal integration of DNA. ACS synthetic biology 2, 537–541 (2013).

25. Tellier, M., Bouuaert, C. C. & Chalmers, R. Mariner and the ITm Superfamily of Transposons. Microbiol Spectr 3, MDNA3–0033–2014. (2015).

26. van Opijnen, T. & Camilli, A. Transposon insertion sequencing: a new tool for systems-level analysis of microorganisms. Nat Rev Microbiol 11, 435–442 (2013).

27. Haniford, D. B. & Ellis, M. J. Transposons Tn10 and Tn5. Microbiol Spectr 3, MDNA3–0002–2014 (2015).

28. Goodall, E. C. A. et al. The Essential Genome of Escherichia coliK-12. mBio 9, 385–18 (2018).

29. Chen, S. P. & Wang, H. H. An Engineered Cas-Transposon System for Programmable and Site-Directed DNA Transpositions. The CRISPR Journal 2, 376–394 (2019).

30. Bhatt, S. & Chalmers, R. Targeted DNA transposition in vitro using a dCas9-transposase fusion protein. Nucleic Acids Res. 6, 7–10 (2019).

31. Enyeart, P. J., Mohr, G., Ellington, A. D. & Lambowitz, A. M. Biotechnological applications of mobile group II introns and their reverse transcriptases: gene targeting, RNA-seq, and non-coding RNA analysis. Mob DNA 5, 2 (2014).

32. Esvelt, K. M. & Wang, H. H. Genome-scale engineering for systems and synthetic biology. Mol Syst Biol 9, 641 (2013).

33. Perutka, J., Wang, W., Goerlitz, D. & Lambowitz, A. M. Use of computer-designed group II introns to disrupt Escherichia coli DExH/D-box protein and DNA helicase genes. J Mol Biol 336, 421–439 (2004).

34. Karberg, M. et al. Group II introns as controllable gene targeting vectors for genetic manipulation of bacteria. Nat Biotechnol 19, 1162–1167 (2001).

35. Klompe, S. E., Vo, P. L. H., Halpin-Healy, T. S. & Sternberg, S. H. Transposon-encoded CRISPR-Cas systems direct RNA-guided DNA integration. Nature 571, 219–225 (2019).

36. Peters, J. E., Makarova, K. S., Shmakov, S. & Koonin, E. V. Recruitment of CRISPR-Cas systems by Tn7-like transposons. Proc Natl Acad Sci USA 114, E7358–E7366 (2017).

37. Halpin-Healy, T. S., Klompe, S. E., Sternberg, S. H. & Fernández, I. S. Structural basis of DNA targeting by a transposon-encoded CRISPR-Cas system. Nature 577, 271–274 (2020).

38. Faure, G. et al. CRISPR-Cas in mobile genetic elements: counter-defence and beyond. Nat Rev Microbiol 17, 513–525 (2019).

39. Peters, J. E. Targeted transposition with Tn7 elements: safe sites, mobile plasmids, CRISPR/Cas and beyond. Mol Microbiol 112, 1635–1644 (2019).

40. Strecker, J. et al. RNA-guided DNA insertion with CRISPR-associated transposases. Science 365, 48–53 (2019).

41. Chavez, M. & Qi, L. S. Site-Programmable Transposition: Shifting the Paradigm for CRISPR-Cas Systems. Mol Cell 75, 206–208 (2019).

42. Hou, Z. & Zhang, Y. Inserting DNA with CRISPR. Science 365, 25–26 (2019).

43. Ronda, C., Chen, S. P., Cabral, V., Yaung, S. J. & Wang, H. H. Metagenomic engineering of the mammalian gut microbiome in situ. Nat Meth 16, 167–170 (2019).

44. Høyland-Kroghsbo, N. M., Muñoz, K. A. & Bassler, B. L. Temperature, by Controlling Growth Rate, Regulates CRISPR-Cas Activity in Pseudomonas aeruginosa. mBio 9, (2018).

45. Stellwagen, A. E. & Craig, N. L. Avoiding self: two Tn7-encoded proteins mediate target immunity in Tn7 transposition. EMBO J 16, 6823–6834 (1997).

46. Greene, E. C. & Mizuuchi, K. Target immunity during Mu DNA transposition. Transpososome assembly and DNA looping enhance MuA-mediated disassembly of the MuB target complex. Mol Cell 10, 1367–1378 (2002).

47. Hagemann, A. T. & Craig, N. L. Tn7 transposition creates a hotspot for homologous recombination at the transposon donor site. Genetics 133, 9–16 (1993).

48. Lin, M. T. et al. Escherichia coli auxotroph host strains for amino acid-selective isotope labeling of recombinant proteins. Meth Enzymol 565, 45–66 (2015).

49. Hickman, A. B. & Dyda, F. DNA Transposition at Work. Chem Rev 116, 12758–12784 (2016).

50. Abbas, A. F., Al-Saadi, A. G. M. & Alkhudhairy, M. K. Biofilm Formation and Virulence Determinants of Klebsiella oxytoca Clinical Isolates from Patients with Colorectal Cancer. J Gastrointest Cancer 72, 2787–6 (2019).

51. Kim, D.-K. et al. Metabolic engineering of a novel Klebsiella oxytoca strain for enhanced 2,3-butanediol production. J. Biosci. Bioeng. 116, 186–192 (2013).

52. Loeschcke, A. & Thies, S. Pseudomonas putida-a versatile host for the production of natural products. Appl Microbiol Biotechnol 99, 6197–6214 (2015).

53. Nikel, P. I. & de Lorenzo, V. Pseudomonas putida as a functional chassis for industrial biocatalysis: From native biochemistry to trans-metabolism. Metab. Eng. 50, 142–155 (2018).

54. Sun, J. et al. Genome editing and transcriptional repression in Pseudomonas putida KT2440 via the type II CRISPR system. Microb. Cell Fact. 17, 41 (2018).

55. Wirth, N. T., Kozaeva, E. & Nikel, P. I. Accelerated genome engineering of Pseudomonas putida by I-SceI-mediated recombination and CRISPR-Cas9 counterselection. Microb Biotechnol 13, 233–249 (2020).

56. Tsai, S. Q. & Joung, J. K. Defining and improving the genome-wide specificities of CRISPR-Cas9 nucleases. Nature reviews Genetics 17, 300–312 (2016).

57. Valderrama, J. A., Kulkarni, S. S., Nizet, V. & Bier, E. A bacterial gene-drive system efficiently edits and inactivates a high copy number antibiotic resistance locus. Nat Commun 10, 5726–8 (2019).

58. Aparicio, T., de Lorenzo, V. & Martínez-García, E. CRISPR/Cas9-enhanced ssDNA recombineering for Pseudomonas putida. Microb Biotechnol (2019). doi:10.1111/1751-7915.13453

59. Duque, E. et al. Identification and elucidation of in vivo function of two alanine racemases from Pseudomonas putida KT2440. Environ Microbiol Rep 9, 581–588 (2017).

60. Langmead, B. & Salzberg, S. L. Fast gapped-read alignment with Bowtie 2. Nat Meth 9, 357–359 (2012).

61. Jung, C. et al. Massively Parallel Biophysical Analysis of CRISPR-Cas Complexes on Next Generation Sequencing Chips. Cell 170, 35–47.e13 (2017).

