## Supplementary Figures for "CRISPR RNA-guided integrases for high-efficiency and multiplexed bacterial genome engineering"

[1]Department of Pharmacology, Columbia University Irving Medical Center, New York, NY, USA. [2]Department of Systems Biology, Columbia University Irving Medical Center, New York, NY, USA. [3]Department of Biochemistry and Molecular Biophysics, Columbia University, New York, NY, USA. [4]Department of Biological Sciences, Columbia University, New York, NY, USA. [5]Department of Pathology and Cell Biology, Columbia University Irving Medical Center, New York, NY, USA.

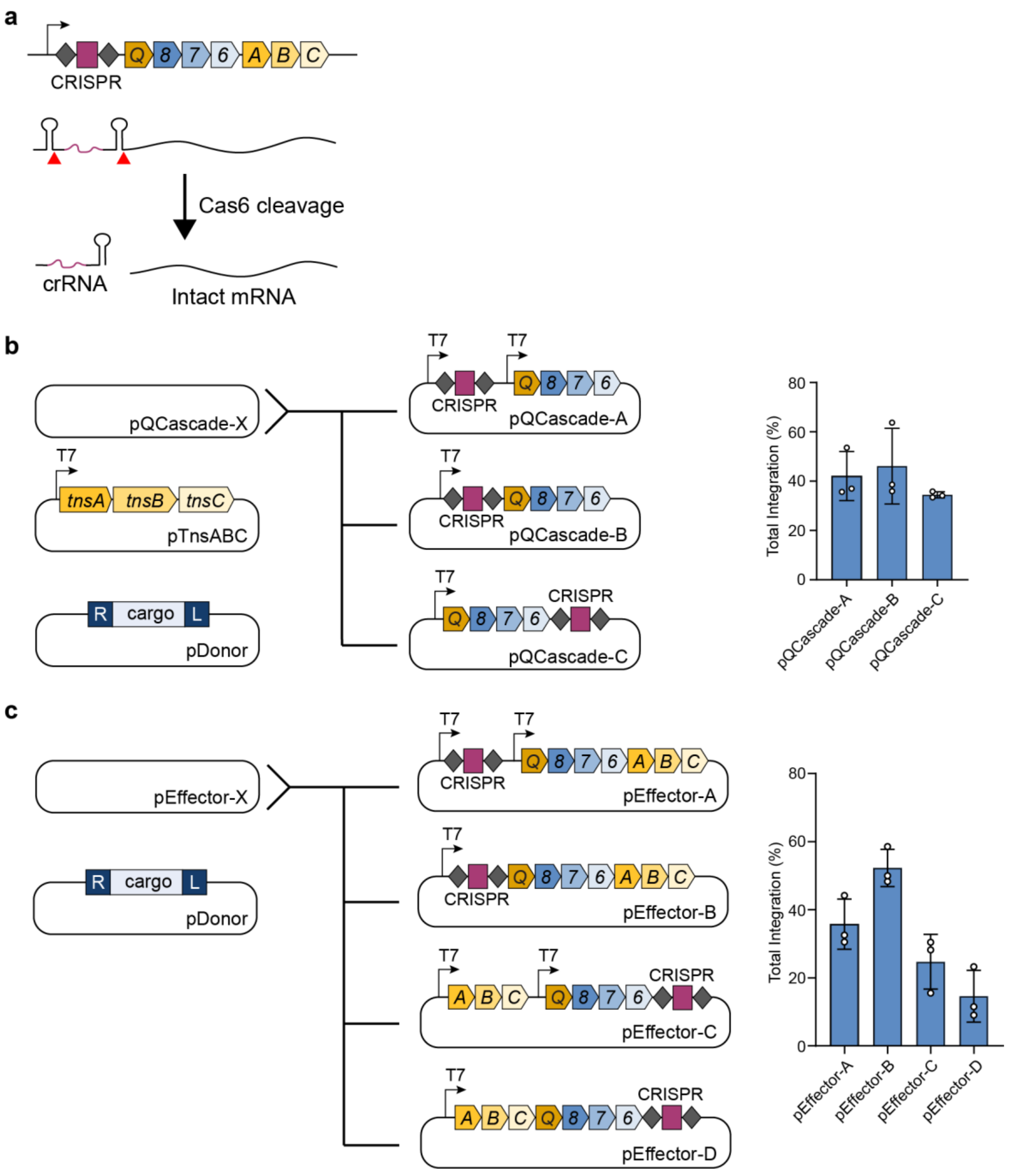


**Supplementary Fig. 1 | Reduction of promoter and plasmid requirements for RNA-guided DNA integration. a,** Schematic illustrating Cas6-dependent processing of an RNA transcript comprising precursor CRISPR RNA and polycistronic mRNA, which liberates the mature crRNA; CRISPR repeats are shown as hairpins. **b,** Left, three pQCascade designs containing either two or one T7 promoters, with the CRISPR array either upstream of downstream of the operon. Right, qPCR-based quantification of integration efficiency with crRNA-4. Cells contained pDonor, pTnsABC, and the indicated pQCascade construct. **c,** Left, four protein-RNA expression plasmid constructs containing either two or one T7 promoters, with the CRISPR array either upstream of downstream of the operon. Right, qPCR-based quantification of integration efficiency with crRNA-4. Cells contained pDonor and the indicated expression plasmid. Data in **b** and **c** are shown as mean ± s.d. for n = 3 biologically independent samples.

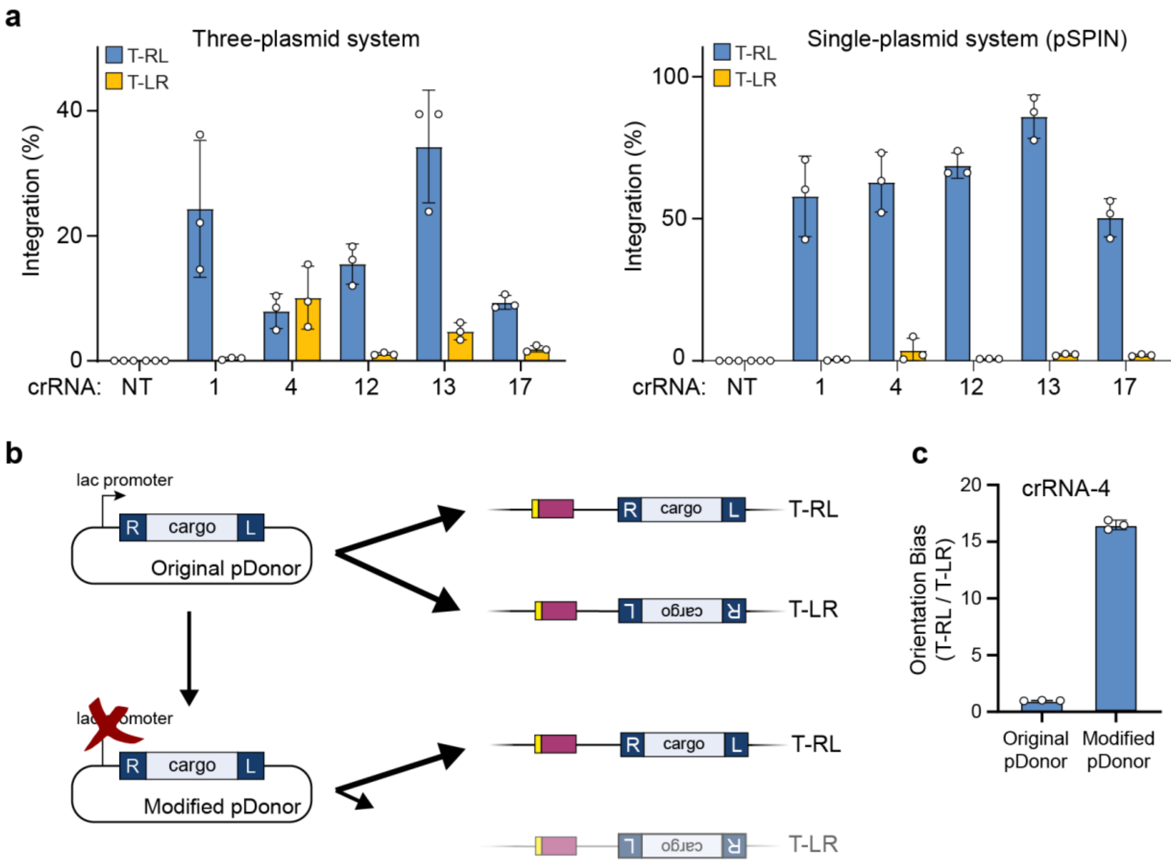


**Supplementary Fig. 2 | Mini-Tn vector context effects on integration orientation. a,** Integration efficiencies in the T-RL and T-LR orientation are plotted from experiments in **Fig. 1e**, for the three-plasmid (left) and single-plasmid (right) expression systems. Integration is more heavily biased towards T-RL for the single-plasmid system, particularly for crRNA-4. **b,** Schematic of the original pDonor plasmid, which contains a lac promoter upstream of the transposon right end, and a modified pDonor plasmid in which this promoter was removed. The modified pDonor shows more frequent T-RL integration, which may be due to the absence of active transcription across the right (R) transposon end. **c,** Comparison of integration orientation bias (T-RL : T-LR) for the three-plasmid expression system with crRNA-4, using the original or modified pDonor; efficiencies were measured by qPCR. Data in **a** and **c** are shown as mean ± s.d. for n = 3 biologically independent samples.

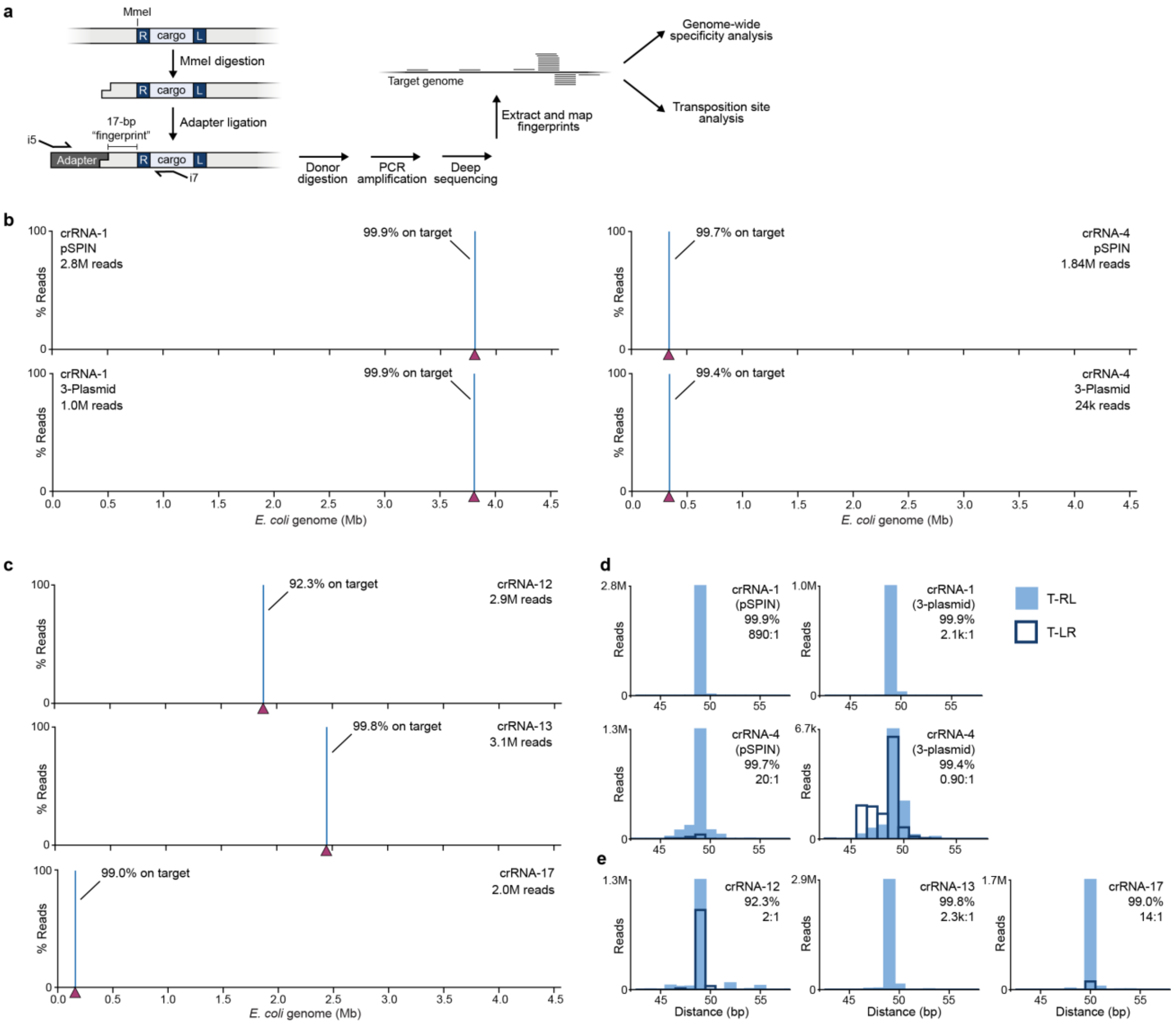


**Supplementary Fig. 3 | Genome-wide analysis of RNA-guided DNA integration by Tn-seq. a,** Tn-seq workflow for deep sequencing of genome-wide transposition events (**Methods**). **b,** Genome-wide distribution of genome-mapping Tn-seq reads for crRNA-1 (left) and crRNA-4 (right) using either the single-plasmid (top) or three-plasmid (bottom) expression system; the target site is denoted by a maroon triangle. **c,** Tn-seq for additional crRNAs using the single-plasmid expression system, shown as in **b**. **d,** Integration site distributions for crRNA-1 (top) and crRNA-4 (bottom) using either the single-plasmid (top) or three-plasmid (bottom) expression system, determined from the Tn-seq data; the distance between the target site and mini-Tn insertion site is shown. Data for both integration orientations are superimposed, with filled blue bars and dark outlines representing T-RL and T-LR, respectively. Values in the top-right corner of each graph give the on-target specificity (%), calculated as the percentage of reads resulting from integration within 100 bp of the primary integration site compared to all genome-mapping reads, and the orientation bias (*X:Y*), calculated as the ratio of T-RL : T-LR reads within the on-target window. **e,** Integration site distributions for additional crRNAs using the single-plasmid expression system, shown as in **d**.

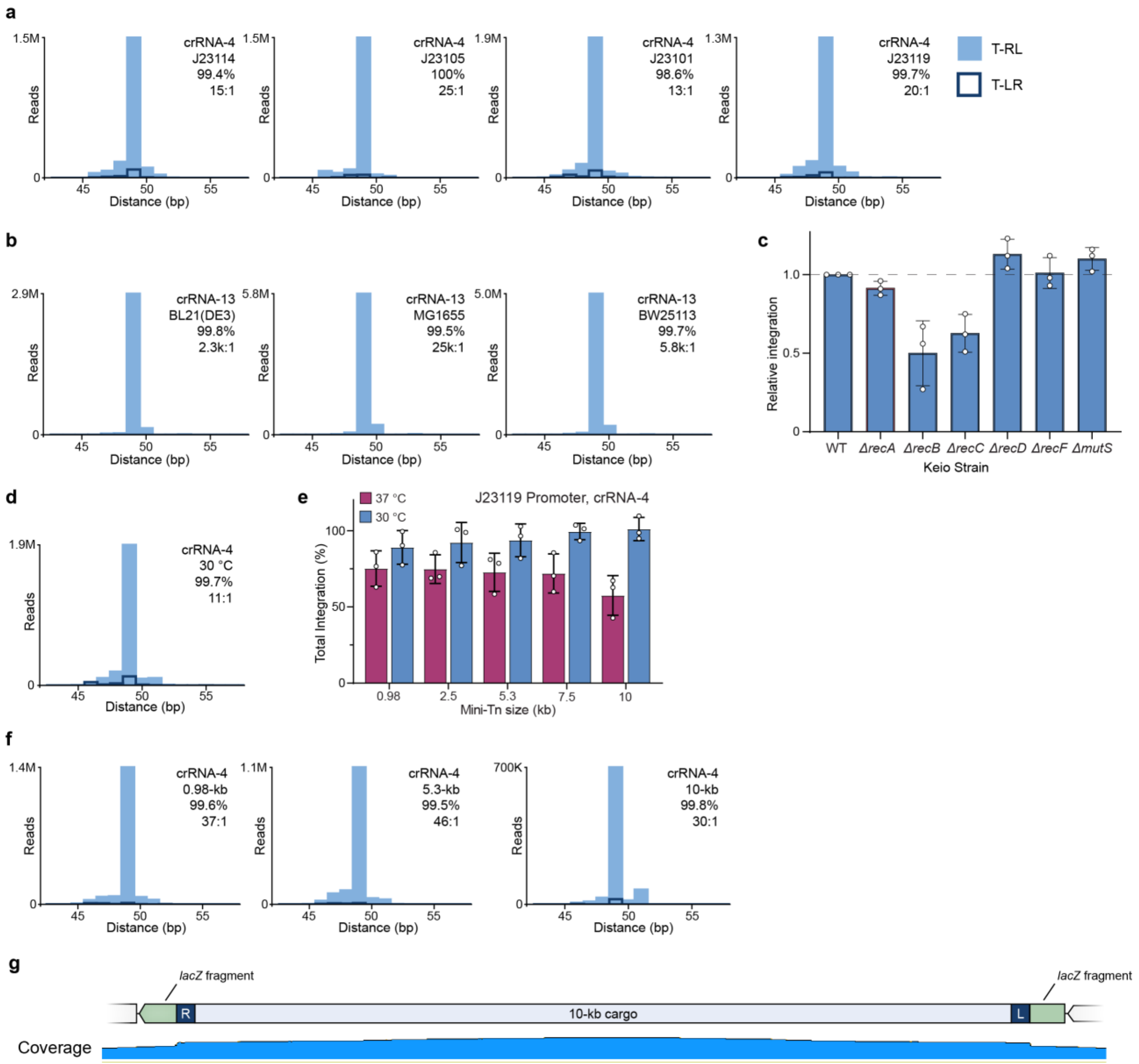


**Supplementary Fig. 4 | Analysis of genome-wide integration specificity as a function of promoter strength, cargo size, and *E. coli* strain. a,** Integration site distributions for crRNA-4 as a function of promoter strength, determined from the Tn-seq data; the distance between the target site and mini-Tn insertion site is shown. Data for both integration orientations are superimposed, with filled blue bars and dark outlines representing T-RL and T-LR, respectively. Values in the top-right corner of each graph give the on-target specificity (%), calculated as the percentage of reads resulting from integration within 100 bp of the primary integration site compared to all genome-mapping reads, and the orientation bias (*X:Y*), calculated as the ratio of T-RL : T-LR reads within the on-target window. **b,** Integration site distributions for crRNA-13, determined for three different laboratory strains of *E. coli*, shown as **a**. **c,** qPCR-based quantification of integration efficiency for crRNA-13 in the indicated Keio knockout strains; integration efficiency was reduced for the *ΔrecB* and *ΔrecC* strains, but unaffected in *ΔrecA*, *ΔrecD*, *ΔrecF*, and *ΔmutS* strains. Data are normalized to the efficiency in the WT BW25113 parental strain. **d,** Integration site distribution for crRNA-4 under control of the J23119 promoter after cells were cultured at 30 °C, shown as in **a**. **e,** qPCR-based quantification of integration efficiency for variable mini-Tn sizes after culturing at either 30 or 37 °C. The promoter and crRNA used are shown at top. **f,** Integration site distributions for crRNA-4 as a function of cargo size, shown as in. **g,** Whole-genome, single-molecule real-time (SMRT) sequencing data for an isolated clone containing the 10-kb insertion, shown as coverage of aligned reads across the entire locus. Data in **c** and **e** are shown as mean ± s.d. for n = 3 biologically independent samples.

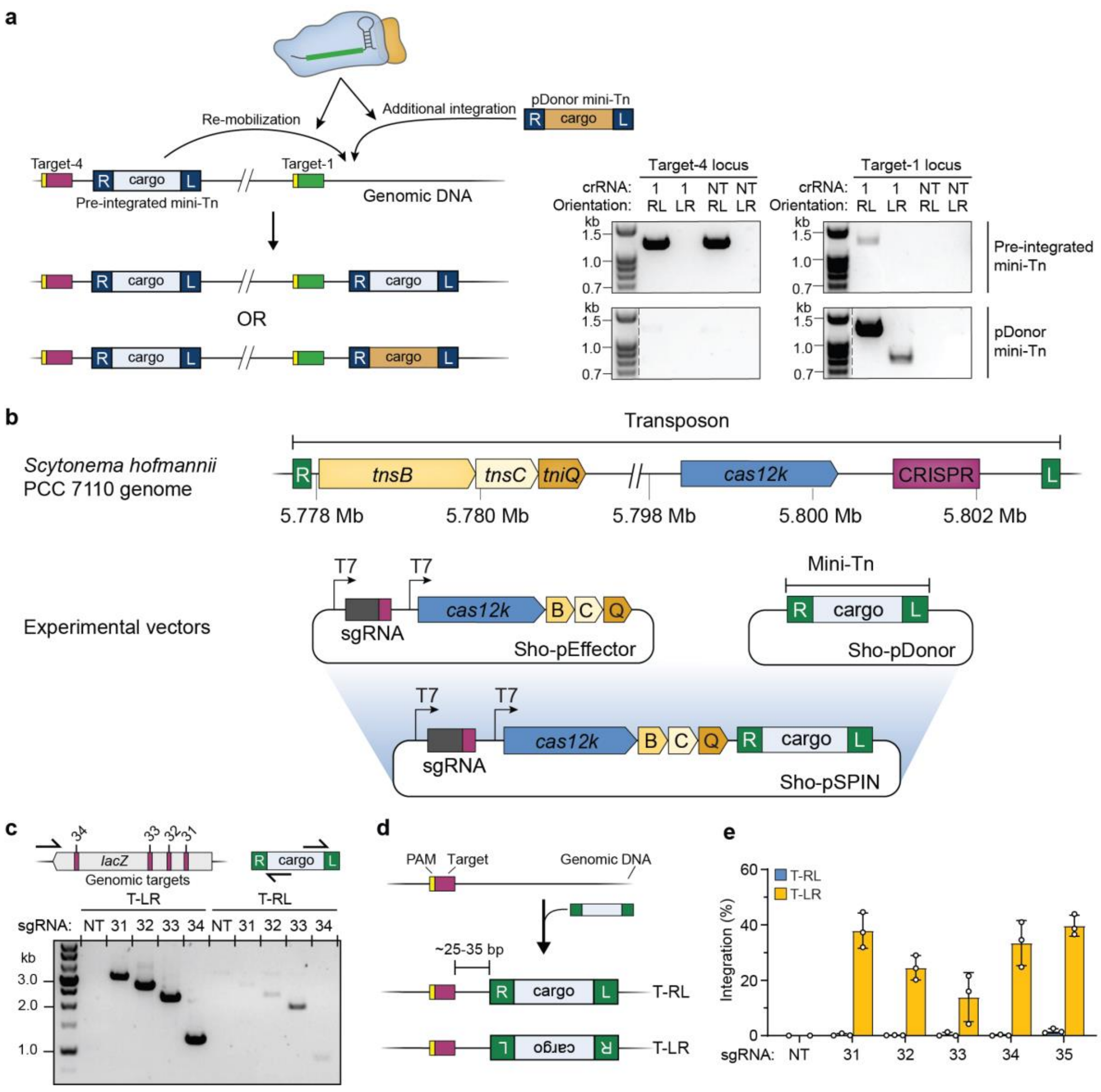


**Supplementary Fig. 5 | Evaluation of mini-Tn remobilization by Vch INTEGRATE, and characterization of a new Type V-K *S. hofmannii* INTEGRATE system. a,** Left, schematic showing potential competition between a genomic- and pDonor-borne mini-Tn when a new site is targeted for RNA-guided DNA integration; the two possible products can be discriminated by cargo-specific primer binding sites. Right, PCR products probing for transposition of the genomic mini-Tn (top) or pDonor-borne mini-Tn (bottom) to the target-1 locus. Although pDonor is the preferred substrate, there is also detectable re-mobilization of the genomic mini-Tn substrate, without apparent loss of the mini-Tn at target-4. **b,** Top, native genomic organization of a Type V-K CRISPR-transposon encoding Cas12k, found within the genome of *Scytonema hofmannii* (Sho) strain PCC 7110; note that this transposon is distinct from that reported elsewhere from the same species[40]. Bottom, plasmid constructs used to recombinantly express the sgRNA and protein components (Sho-pGCT) and the mini-Tn (Sho-pDonor). **c,** Genomic locus targeted by sgRNAs 31–34 (top), and PCR analysis of transposition by ShoINT, resolved by agarose gel electrophoresis (bottom). Bidirectional integration was observed in both T-RL and T-LR orientations for multiple sgRNAs, though there is a strong bias for T-LR. **d,** Overview of RNA-guided DNA integration by ShoINT. Insertion occurs in two possible orientations, similarly to the Type I-F VchINT system, at an approximate distance of 25-35 bp from the edge of the target site. The 4-nt PAM and 23-nt protospacer are shown as orange and maroon rectangles, respectively. **e,** qPCR-based quantification of integration efficiency for sgRNAs 31–35. Data in **e** are shown as mean ± s.d. for n = 3 biologically independent samples.

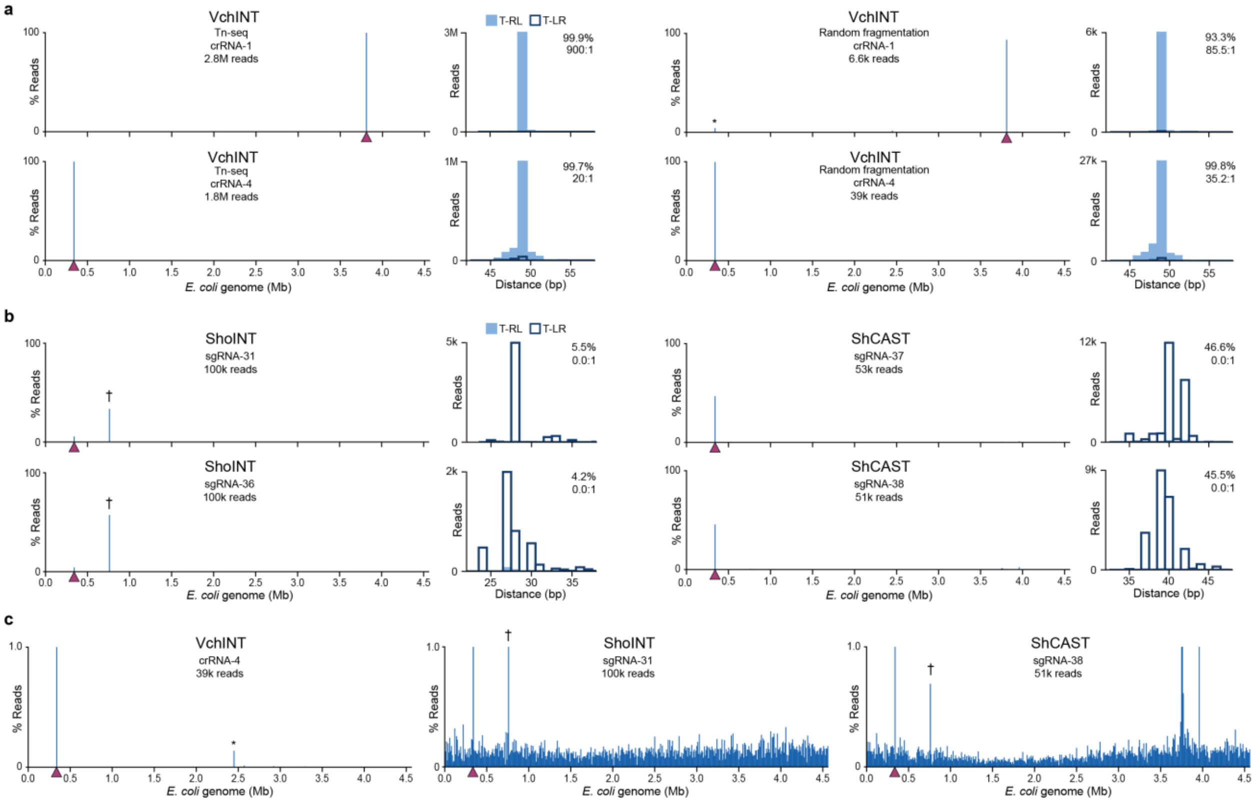


**Supplementary Fig. 6 | Analysis of genome-wide integration events for three CRISPR-transposon systems. a,** Comparison of two distinct next-generation sequencing (NGS) library preparation techniques for analyses of genome-wide integration specificity with VchINT: transposon-insertion sequencing (Tn-seq; left), based on restriction digestion and adaptor ligation onto mini-Tn-containing genomic fragments, followed by targeted PCR; and random fragmentation (right) and adaptor ligation onto all genomic fragments, followed by targeted PCR. The target site is denoted by a maroon triangle. Insets show integration site distributions determined from the NGS data; the distance between the target site and mini-Tn insertion site is shown. Data for both integration orientations are superimposed, with filled blue bars and dark outlines representing T-RL and T-LR, respectively. Values in the top-right corner of each graph give the on-target specificity (%), calculated as the percentage of reads resulting from integration within 100 bp of the primary integration site compared to all genome-mapping reads, and the orientation bias (*X:Y*), calculated as the ratio of T-RL : T-LR reads within the on-target window. Both analyses return highly consistent data. **b,** Analysis of genome-wide integration specificity with ShoINT (left) and the ShCAST system described previously[40] (right), shown as in **a**. ShoINT exhibited high levels of integration into the T7 RNAP gene (†), suggesting a cellular fitness benefit when expression of the recombinant protein-RNA machinery is eliminated through T7 RNAP inactivation. **c,** Comparison of genome-wide specificity between VchINT (Type I-F), ShoINT (Type V-K), and ShCAST (Type V-K) as assessed via random fragmentation-based NGS library preparation, shown as in **a** but focused on reads comprising 1% or less of the library. The Type I-F system exhibits exquisite accuracy, whereas both Type V-K systems exhibit rampant non-specific integration across the *E. coli* genome. *, low-level, well-to-well contamination of NGS data from other samples.

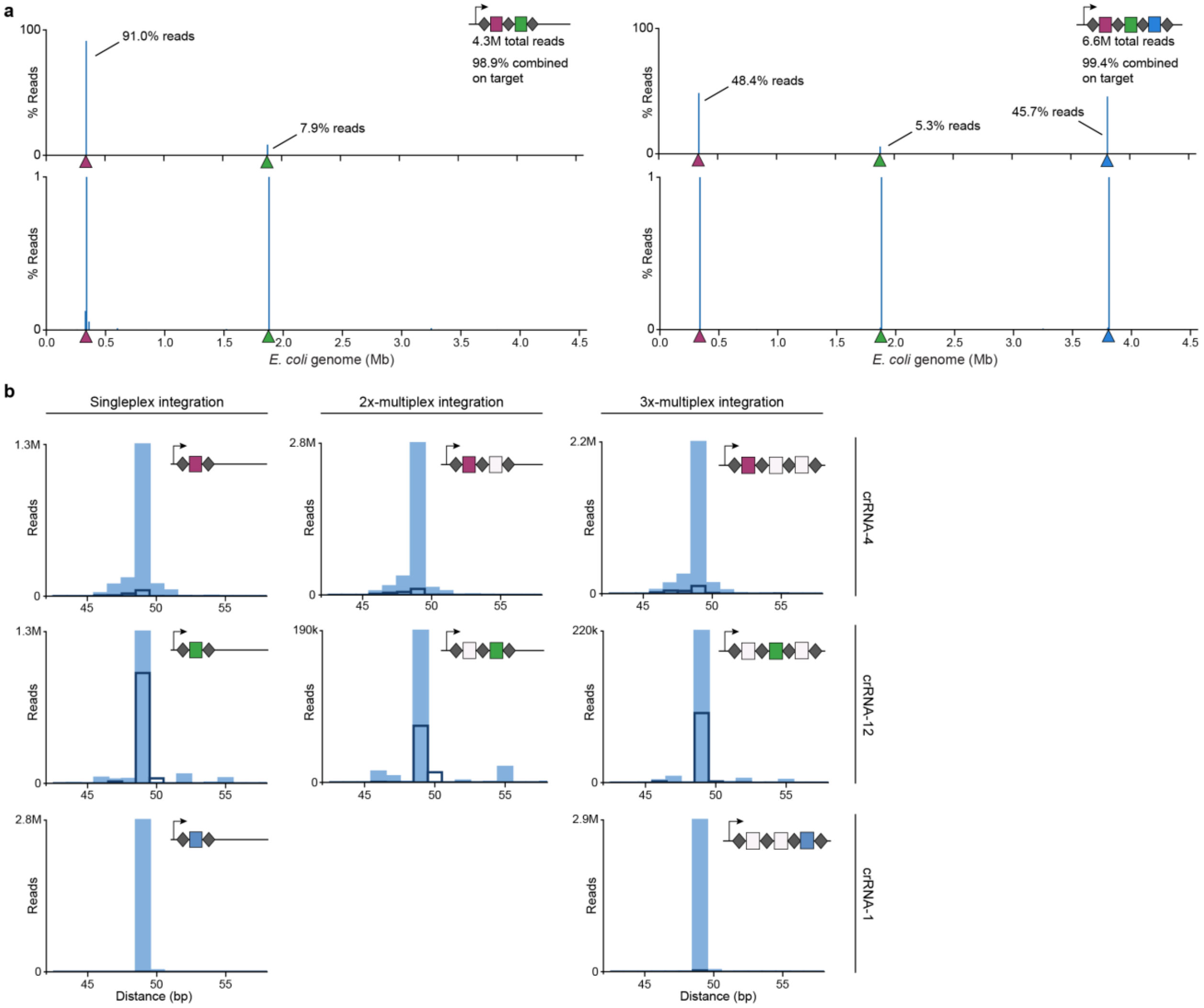


**Supplementary Fig. 7 | Genome-wide analysis of multiplexed RNA-guided DNA integration. a,** Genome-wide distribution of genome-mapping Tn-seq reads for a double-spacer (left) and triple-spacer (right) CRISPR array; the corresponding target sites are denoted by similarly colored triangles. The top graphs plot the percentage of total reads; the bottom graphs focus on reads comprising 1% or less of the library, revealing an absence of detectable off-target events. The overall on-target percentages combine all reads mapping to the on-target window of each individual genomic target. **b,** Integration site distributions for the indicated crRNA as a function of CRISPR array composition, determined from the Tn-seq data; the distance between the target site and mini-Tn insertion site is shown. Data for both integration orientations are superimposed, with filled blue bars and dark outlines representing T-RL and T-LR, respectively.

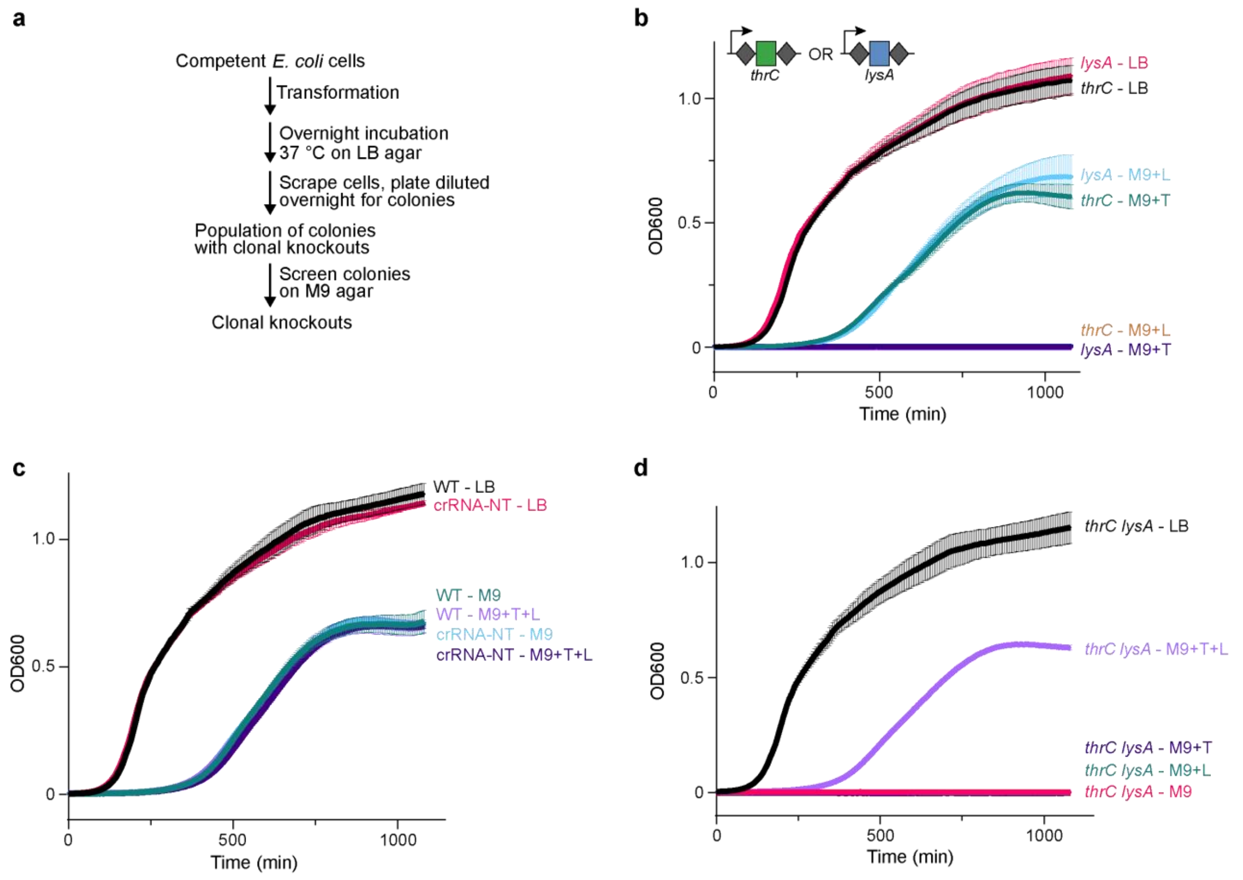


**Supplementary Fig. 8 | Generation of auxotrophic *E. coli* strains through single- or multiplex integration. a,** Workflow for generating and screening auxotrophic *E. coli* knockouts with multiplexed RNA-guided DNA integration (**Methods**). **b**, Growth curves for single-knockout *E. coli* clones cultured at 37 °C in LB or M9 minimal media with or without supplemented threonine (T) and lysine (L). **c**, Growth curves for WT or control *E. coli* clones transformed with a non-targeting crRNA (crRNA-NT), cultured at 37 °C in LB or M9 minimal media with or without supplemented threonine (T) and lysine (L). **d,** Growth curves for double-knockout *E. coli* clone cultured at 37 °C in LB or M9 minimal media with or without supplemented threonine (T) and lysine (L), after five cycles of serial passaging and overnight growth in LB media. Data in **b**, **c** and **d** are shown as mean ± s.d. for three technical replicates.

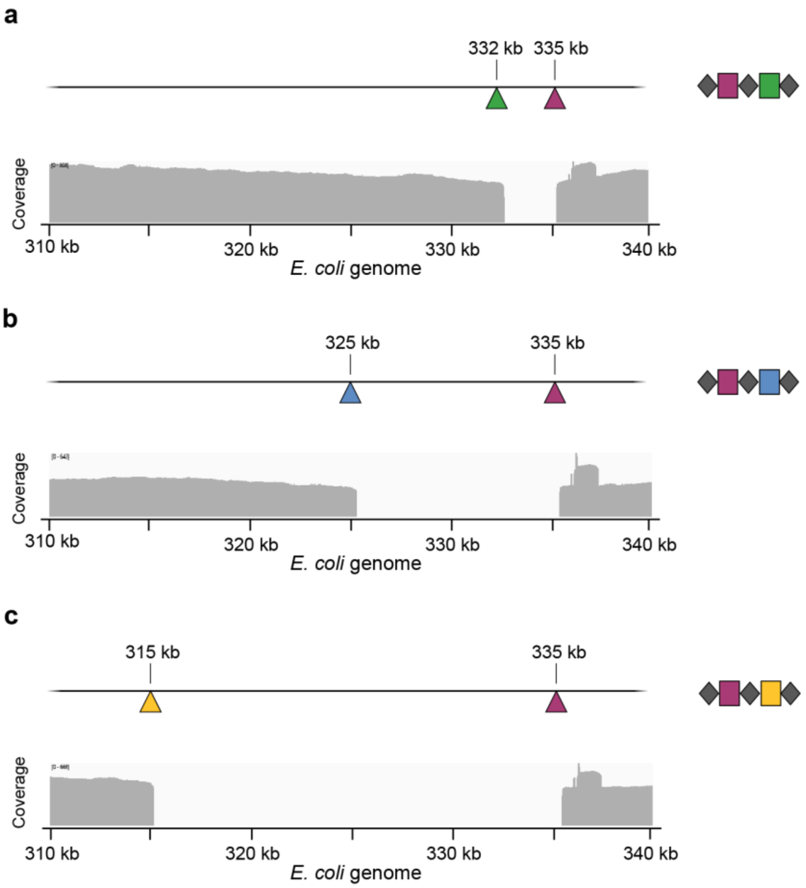


**Supplementary Fig. 9 | SMRT sequencing of programmed deletions using INTEGRATE and Cre-Lox. a,** Top, schematic of genomic locus targeted for a 2.4-kb deletion with the double-spacer CRISPR array shown at the right; triangles represent corresponding target sites. Bottom, coverage data from whole-genome SMRT sequencing reads from an isolated clone, aligned to the *E. coli* BL21(DE3) reference genome. **b,** 10-kb deletion data, shown as in **a**. **c,** 20-kb deletion data, shown as in **a**.

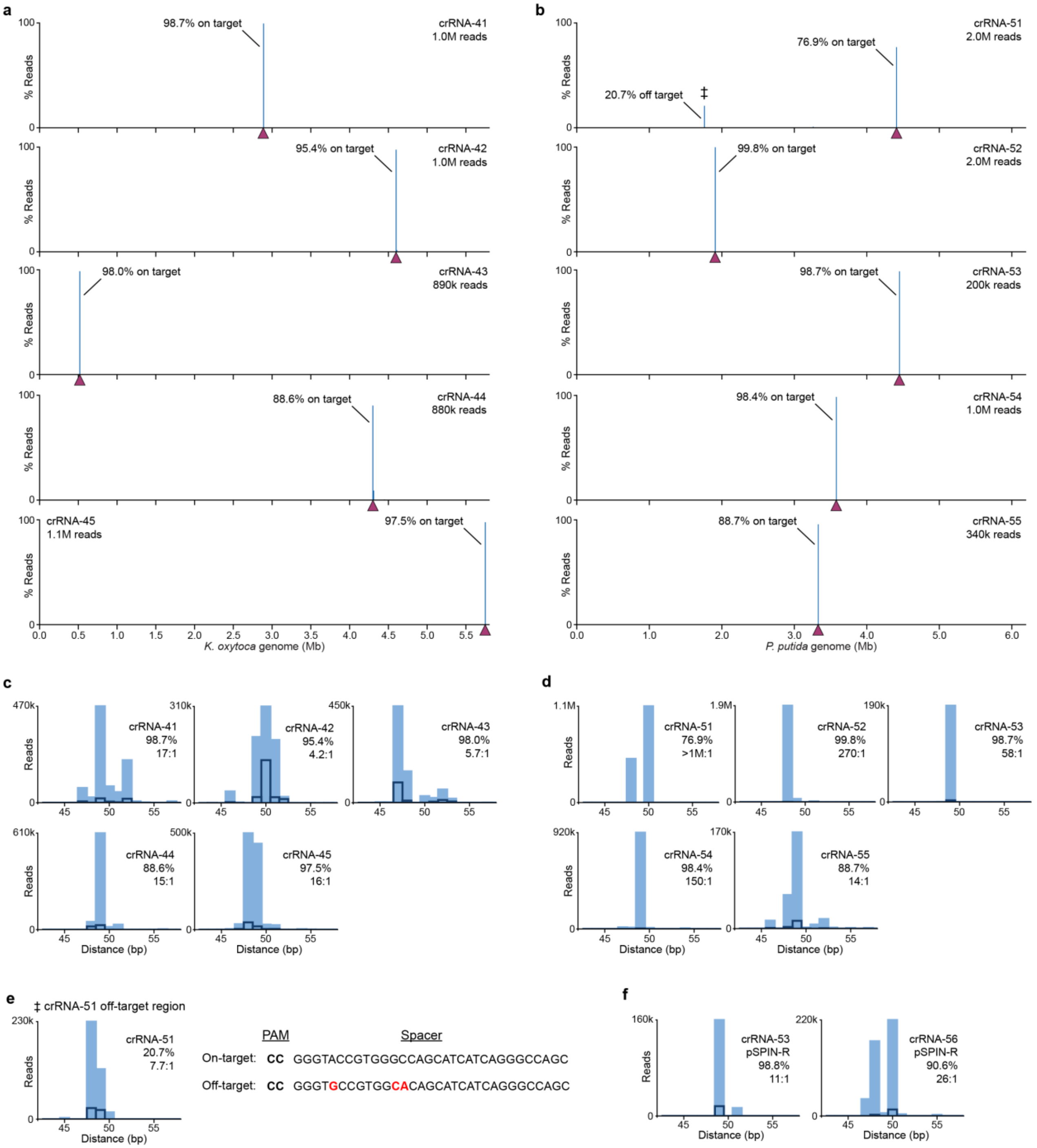


**Supplementary Fig. 10 | Genome-wide analysis of RNA-guided DNA integration in *K. oxytoca* and *P. putida*. a,** Genome-wide distribution of genome-mapping Tn-seq reads for the indicated crRNA expressed by pSPIN-BBR1 in *K. oxytoca*; the target site is denoted by a maroon triangle. **b,** Genome-wide distribution of genome-mapping Tn-seq reads for the indicated crRNA expressed by pSPIN-BBR1 in *P. putida*; the target site is denoted by a maroon triangle. ‡, off-target integration site (see **e**). **c,** Integration site distributions for the indicated crRNAs in *K. oxytoca*, determined from the Tn-seq data; the distance between the target site and mini-Tn insertion site is shown. Data for both integration orientations are superimposed, with filled blue bars and dark outlines representing T-RL and T-LR, respectively. Values in the top-right corner of each graph give the on-target specificity (%), calculated as the percentage of reads resulting from integration within 100 bp of the primary integration site compared to all genome-mapping reads, and the orientation bias (*X:Y*), calculated as the ratio of T-RL : T-LR reads within the on-target window. **d,** Integration site distributions for the indicated crRNAs in *P. putida*, shown as in **c**. **e,** Integration site distributions for the off-target peak (‡) with crRNA-51 in *P. putida*, shown in **c**. The sequences of the on-target and off-target sequences upstream of the integration site are shown to the right, highlighting the high degree of sequence similarity. **f,** Integration site distributions for the indicated crRNAs in *P. putida*, shown as in **d**; these experiments utilized the reversed pSPIN-R plasmid, as compared to the pSPIN plasmid used in **d**.

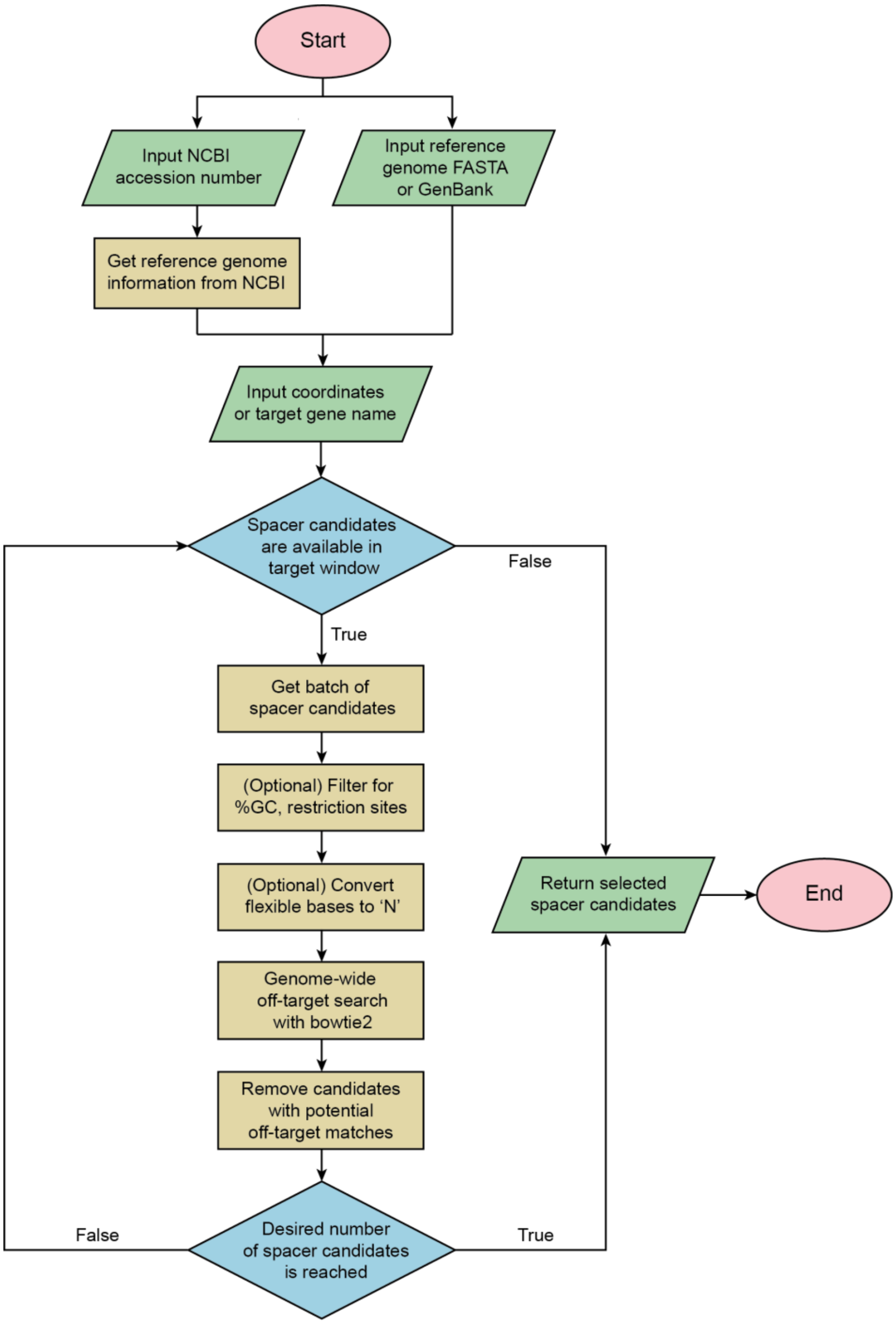


**Supplementary Fig. 11 | Flowchart for the INTEGRATE guide RNA design algorithm.** Spacers with a defined length and PAM are generated and filtered from a given reference genome, based on the target gene name or genomic coordinates. The Bowtie2 alignment tool[60] is used to evaluate each spacer candidate for potential off-targets genome-wide. Spacers are considered to have potential off-targets when Bowtie2 detects alignments exhibiting less than a user-specified maximum mismatch limit. For bacterial genomes, we find that this process usually results in a sufficient number of spacers within each window, without the need for scoring each spacer candidate. For Type I Cascade (such as VchINT) spacers, the program converts flexible bases – those bases occurring every 6$^{th}$ position, which do not contribute to spacer-protospacer complementarity within the R-loop[37,61] – to 'N' to exclude these bases from contributing to the mismatch count for the genome-wide off-target search. The off-target search module can also be executed separately for the evaluation of user-specified spacers. The program and more in-depth documentation are publicly accessible via GitHub (https://github.com/sternberglab/INTEGRATE-guide-RNA-tool).
